# Distinguishing between recent balancing selection and incomplete sweep using deep neural networks

**DOI:** 10.1101/2020.07.31.230706

**Authors:** Ulas Isildak, Alessandro Stella, Matteo Fumagalli

## Abstract

Balancing selection is an important adaptive mechanism underpinning a wide range of phenotypes. Despite its relevance, the detection of recent balancing selection from genomic data is challenging as its signatures are qualitatively similar to those left by ongoing positive selection. In this study we developed and implemented two deep neural networks and tested their performance to predict loci under recent selection, either due to balancing selection or incomplete sweep, from population genomic data. Specifically, we generated forward-in-time simulations to train and test an artificial neural network (ANN) and a convolutional neural network (CNN). ANN received as input multiple summary statistics calculated on the locus of interest, while CNN was applied directly on the matrix of haplotypes. We found that both architectures have high accuracy to identify loci under recent selection. CNN generally outperformed ANN to distinguish between signals of balancing selection and incomplete sweep and was less affected by incorrect training data. We deployed both trained networks on neutral genomic regions in European populations and demonstrated a lower false positive rate for CNN than ANN. We finally deployed CNN within the *MEFV* gene region and identified several common variants predicted to be under incomplete sweep in a European population. Notably, two of these variants are functional changes and could modulate susceptibility to Familial Mediterranean Fever, possibly as a consequence of past adaptation to pathogens. In conclusion, deep neural networks were able to characterise signals of selection on intermediate-frequency variants, an analysis currently inaccessible by commonly used strategies.

## 2 Introduction

Balancing selection is a selective process that generates and maintains genetic diversity within populations, as firstly proposed by Dobzhansky in 1951 [1]. Many diverse mechanisms of balancing selection have been described [2]. Over-dominance (or heterozygote advantage) occurs when heterozygote individuals at one locus have higher fitness than homozygotes. In sexually antagonistic selection, different alleles at the same locus have opposite effects in the two sexes creating a balanced polymorphism at the population level. In negative frequency-dependent selection, rare alleles have a fitness advantage. Finally, spatially and temporally varying selection creates a scenario where different alleles are advantageous in different environments.

Until 2006 the general consensus was that only few loci in the human genome have been targets of balancing selection [3, 4]. Since then, the availability of large-scale population genomics data and the development of *ad hoc* statistical tests contributed to the current view that balancing selection is a widespread adaptive mechanism underlying a broad spectrum of features in the genetic architecture of phenotypes [5, 6].

In humans, balancing selection is responsible for shaping the diversity of genes involved in the adaptive and innate immune response [7, 8, 9, 10], metabolism [11] and other processes [12]. Notably, variants targeted by pathogen-driven balancing selection have been found to be associated with susceptibility to several autoimmune diseases [13]. Therefore, by elucidating the genomic signals of balancing selection we have the ability to identify common alleles with critical functional consequences. For instance, balancing selection has been hypothesised to maintain a common variant in an angiotensin-converting enzyme [14] which has been recently associated to increased susceptibility to SARS-CoV-2 [15].

Several methods to identify targets of balancing selection have been proposed [16]. Genomic signatures of balancing selection have been detected by testing for an excess of heterozygous genotypes [17], a local increase in genetic diversity [18] and polymorphisms [19], a shift in the site frequency spectrum towards common frequencies [9, 20, 12], a population genetic differentiation lower or higher than expected under neutral evolution [21], presence of trans-species polymorphism [22, 23], by explicitly modelling the patterns of polymorphisms and substitutions [10, 24], and by correlating allele frequencies with environmental variables [25].

The application of such methods to large-scale human population genomic data has enabled the characterisation of targets of long-term balancing selection (i.e. selection that predates the time to the most recent common ancestor in a species) in humans and their association to several diseases [20, 12]. Nevertheless, all these studies contributed little to the understanding of the role of balancing selection in recent human evolution, despite short-term or transient balancing selection being predicted to be a common phenomenon in nature [26]. Recent balancing selection leaves traces that are almost indistinguishable from those left by recent positive selection [16], with beneficial alleles segregating at intermediate frequency in contemporary genomes in both cases [2]. Additionally, even when signatures of balancing selection are identified, the underlying evolutionary mechanism (e.g., overdominance or negative frequency-dependent selection) is often unknown [6]. As such, current methods have only limited power to identify and characterise signatures of recent balancing selection in the human genome.

A promising solution to address this issue is provided by supervised machine learning (ML) which has been recently introduced in population genetics and successfully applied for evolutionary inferences [27]. For instance, several ML methods have been proposed and successfully applied to population genetic data to predict and classify neutral and selective events on genomic loci [28, 29, 30, 31, 32, 33, 34]. Deep learning is a class of ML algorithms based on artificial neural networks (ANNs) which comprise nodes in multiple layers connecting features (input) and responses (output) [35]. ANNs have the potential to be used in population genetics to estimate parameters from genomic data using multiple summary statistics as input [36].

Notably, deep learning algorithms can effectively learn which features (i.e. measurable properties of the data) are sufficient for the prediction [35, 37]. Despite deep learning in population genetics being in its infancy, several studies have already introduced the use of Convolutional Neural Networks (CNNs) to full population genomic data with convolutional layers automatically extracting informative features [38, 39, 40, 41, 42]. A convolution layer is comprised of several weight matrices that slide across the input image and perform a matrix convolutional to produce image matrices [43, 44]. Recent reviews provide more detailed information on convolutional neutral networks in population genetic inference [38, 41].

In this study we aimed at developing and implementing deep neural networks to predict loci at intermediate allele frequency (i.e. between 40% and 60%) under natural selection (Test 1). By doing so, our goal is also to distinguish between signals of incomplete sweep (e.g., ongoing positive selection) and signals of balancing selection (Test 2), either due to overdominance or negative frequency-dependent selection. As mentioned above, these two types of selection are different biologically but leave similar signatures in genomes, making their discernment particularly challenging. Specifically, we compared the predictive power between ANNs (i.e. based on summary statistics) and CNNs (i.e. based on full population genomic data) to perform such classification.

Finally, we deployed the trained deep neural networks on population genomic data to identify and characterise signals of natural selection acting on the *MEFV* gene. Mutations in the *MEFV* gene cause Familial Mediterranean Fever (FMF), an autoinflammatory disease with recurrent episodes of fever, abdominal, joint and chest pain, with gradual development of nephropathic amyloidosis (kidney failure) in some cases [45]. FMF is highly prevalent in populations of Mediterranean origin [45] and the 3’ terminal region of the *MEFV* gene has been hypothesised to be under balancing selection due to overdominance in some European populations [17]. Recently, causative mutations in the *MEFV* gene have been reported as target of recent positive selection in the Turkish population as they confer resistance to *Yersinia pestis* [46]. By applying our deep neural networks on a large sample size of genomic data we sought to establish which type of natural selection has been acting on *MEFV* with regards to susceptibility to FMF.

## 3 Materials and Methods

### 3.1 Simulations of population genomic data

We performed extensive simulations both to assess the predictive power of summary statistics and to train deep neural networks. We generated synthetic population genomic data using *SLiM 3.2*, a forward-in-time genetic simulation software [47]. We simulated four different scenarios: neutrality (NE), incomplete sweep (IS), overdominance (OD) and negative frequency-dependent selection (FD). A locus under balancing selection (BS) was considered to be under either OD or FD. All simulations were conditioned on a previously proposed demographic model for European populations [48] with a mutation rate of 1.44*e −* 8, a generation time of 29 years, and a recombination rate sampled from a Normal distribution with mean 1*e −* 8 and standard deviation 1*e −* 9. Further details on the simulation model employed are available in Table S1.

For simulating scenarios of natural selection, we generated loci of 50k bp (base pairs) with the selected variant at the centre of the simulated sequence. We assumed a model of selection on a *de novo* mutation. For illustrative purposes of this study, the selected mutation was introduced in the European population at 21 different times, ranging from 40k to 20k ya (Figure S1). We classified these times into three categories: recent (20k to 26k ya), medium (27k to 33k ya), and old (34k to 40k ya) selection.

To mimic the effect of a selected variant at intermediate frequency, we conditioned the final (i.e. contemporary) allele frequencies to be between 40% and 60% in the sample. If the final frequency of the selected allele was not within this range, the simulation restarted at the generation where the selected variant was introduced. For each selection scenario and time of onset of selection, we chose selection coefficients and parameters which maximised the probability of the final allele frequency being between 40% and 60% (Table S2). At the end of the simulations, we sampled 198 chromosomes (i.e. haploid individuals) to match the sample size of CEU (Central European) individuals in the 1000 Genomes Project [49].

In the neutral scenario, no selected variant was introduced. Instead, we generated data with a neutral variant at the centre of the sequence with a frequency between 40% and 60%. To achieve this, we (i) simulated a larger region of 500k bp under neutral evolution, (ii) sampled 198 chromosomes, (iii) identified a variant with a frequency between 40% and 60%, (iv) trimmed the large region to obtain a 50k bp locus (Figure S2).

### 3.2 Calculation of summary statistics and genomic images

We processed the simulated genomic data to be received as input to deep neural networks (i.e. both ANN and CNN). For ANN, we summarised each genomic sequence as a vector of all potentially informative summary statistics. Additionally, we divided each simulated 50k bp sequence into two sub-regions: (1) proximal to the selection site (20-30k bp), and (2) distal from the selected site (0-20k bp + 30-50k bp) (Figure S3), similarly to previous studies [50, 36]. For each region, we calculated 32 summary statistics. The main statistics are: nucleotide diversity *π* [51], Watterson’s estimator *θ* [52], Tajima’s *D* [53], linkage disequilibrium (LD) *r*^2^ [54], Kelly’s *Z_nS_* [55], Fu and Li’s *F\** and *D\** [56], H1, H12, H123, H2/H1 [57], iHS [58], EHH [59], Zeng et al.’s *E* [60], Fay and Wu’s *H* [61], *nS_L_* [62], *NCD1* [12], raggedness index [63], observed and expected heterozygosity, haplotype diversity, number of unique haplotypes, and number of singletons. Finally, we included some derivatives of these main statistics, such as mean, median and maximum values of mean pairwise distances calculated for all chromosome pairs in a simulation (Figure S3, Table S3). All summary statistics were calculated using *scikit-allel* library (https://github.com/cggh/scikit-allel) and then scaled using the *StandardScaler* function from *sklearn* library [64]. All scaled summary statistics were considered as input features to the ANN.

For CNN, we created images from the alignment of sampled haplotypes, similar to previous studies [39, 38, 40]. In this data representation, each row of the image is a sampled haplotype (i.e. individual chromosome) and each column corresponds to a specific segregating site. The colour coding indicates if a variant is derived or ancestral, or any other polarisation of alleles (e.g., major/minor, reference/alternate). To disentangle the effect of random sorting of sampled haplotypes [40], we reordered rows of images as follows: (i) sampled haplotypes are divided into two groups based on the presence or absence of the targeted allele, (ii) haplotypes within each of the two groups are sorted separately based on haplotype frequency, (iii) the two sorted groups are combined to obtain the final reordered image. Lastly, to take into account the different dimensions of simulated loci, we resized images into 128 *×* 128 pixels [40] using the *Image* module from *Pillow* package (https://pypi.org/project/Pillow).

### 3.3 Implementation and training of neural networks

Both ANN and CNN models were implemented in Python using *Keras* library with *Tensorflow* backend [65]. ANN model comprises one input, three hidden, and one output fully-connected (i.e. dense) layers. Similar to a previous study [36], the hidden layers consist of 20, 20, and 10 neurons, respectively, all with a Rectified Linear Units (ReLU) activation function. The output layer, which performs the binary classification, consists of a single neuron with a sigmoid (i.e. logistic) activation function. To control for overfitting, in addition to batch normalisation, we used a dropout rate of 0.5 and L2 weight decay of 0.005 across all but the output layers. Models were optimised using the Adam optimizer with a batch size of 64 and a learning rate of 0.005 [66, 67].

The CNN model consisted of three sets of 2D convolution layers, each followed by a batch normalisation layer and ReLU activation layer. A max-pooling layer was also applied after the first two convolution layers. All convolutional layers consisting of 32 filters had a kernel size of 3x3, applied at stride 1. The size of the pooling layers was 2x2, which were applied at stride 2. The convolutional layers were followed by a flatten layer, which transforms a two-dimensional feature matrix into a vector. Finally, we used a fully-connected layer consisting of 128 units that uses the flattened feature vector as an input, followed by an output layer. Again, we used ReLU activation function on the output from the fully connected layer and the sigmoid function for the output layer. We performed extensive hyper-parameter tuning on training data over 25 epochs to optimise values of learning rate (Figure S4), number of units per layer (Figure S5), L2 regularisation (Figure S6), dropout rates (Figure S7), batch normalisation (Figure S8), image reshaping (Figure S9), to maximise accuracy for predicting loci under incomplete sweep or balancing selection (Test 2). A complete list of all hyper-parameter values used in the CNN model is available in Table S4. Further, we performed data augmentation during the training of CNN models by randomly flipping images horizontally (Figure S10) using the *ImageDataGenerator* function from *Keras* [65]. Similarly, we performed hyper-parameter tuning for ANN on 40k training samples over 25 epochs to optimise values of learning rate (Figure S11), dropout and L2 weight decay (Figure S12), scaling (Figure S13), architecture (Figure S14), to maximise accuracy for predicting loci under incomplete sweep or balancing selection (Test 2).

We performed 480, 000 simulations in total for training all deep neural networks. Each single model employed 80, 000 simulated data samples, 64, 000 of them for training and the remaining 16, 000 for validation. All models were trained for 50 epochs each. Testing was performed on approximately 16, 000 data samples. We trained both ANN and CNN to perform two classification tasks: predict loci under natural selection *vs.* neutral evolution (Test 1) and predict loci under balancing selection *vs.* incomplete sweep (Test 2). The predictive power of ANN and CNN for each test was quantified with a confusion matrix, where each row represents the instances of true class and each column the corresponding number of predicted instances.

### 3.4 Prediction of natural selection from genomic data

We deployed the trained networks on phased population genomic data from the 1000 Genomes Project for the CEU population [49]. We filtered all non-biallelic positions and selected all variants with a frequency between 40% and 60% in CEU populations within the *MEFV* gene region. We retrieved 41 such variants and, for each one, generated a haplotype matrix [40] of 50k bp surrounding the putative target variant. We calculated summary statistics (for ANN) and generated images (for CNN) for each variant by applying the same pipeline used for training the networks. Test 2 was performed only on variants predicted to be under selection for Test 1. Genomic annotations were obtained using the *EnsDb.Hsapiens.v75 package* in R [68] and *Gviz* package was used for visualization [69]. We also employed the same procedure on data from 99 randomly sampled individuals of Tuscans in Italy (TSI) from 1000 Genomes Project [49].

We further deployed the trained networks on genomic regions hypothesised to be neutrally evolving. We extracted two putative neutral regions (chr16: 62,852,764 - 62,944,210 and chr16: 63,651,950 - 63,684,341) predicted by the NRE Tool [70] which was run with default parameters for a large region proximal to *MEFV* gene on chromosome 16. We identified a total of 42 biallelic variants with intermediate allele frequency and applied the same procedure aforemen-tioned to predict signals of selection using both trained networks.

### 3.5 Software availability

A Python package called *BaSe* (Balancing Selection) that implements deep neural networks (both ANN and CNN) for the detection of selection and for discerning between incomplete sweep and balancing selection is available at https://github.com/ulasisik/balancing-selection. Data visualizations were performed in R, using *ggplot2* [71], *ggpubr* [72], and *pheatmap* [73] libraries. All remaining analyses were performed in Python.

## 4 Results

### 4.1 Summary statistics are not sufficient to discriminate between balancing selection and incomplete sweep

Our first aim was to test whether commonly used summary statistics were suficient to discriminate between loci under neutrality and natural selection, the latter comprising both incomplete sweep and balancing selection (Test 1). We calculated a total of 64 different summary statistics and compared their distributions calculated on simulated loci under either neutrality or selection, with the targeted allele at intermediate frequency (between 40% and 60%) in the centre of the region (Figure S15). Figure 1 (upper panel a) shows a subset of these comparisons and indicates that the distribution of several summary statistics under neutral evolution or natural selection are statistically different. Therefore, these summary statistics can be used to predict loci under natural selection. This effect is particularly notable for haplotype-based summary statistics (Figure 1, upper left panel a) and it is consistent across all times of onset of selection (recent, medium, old), in line with the effect of recent selection on patterns of LD.

**Figure 1:**
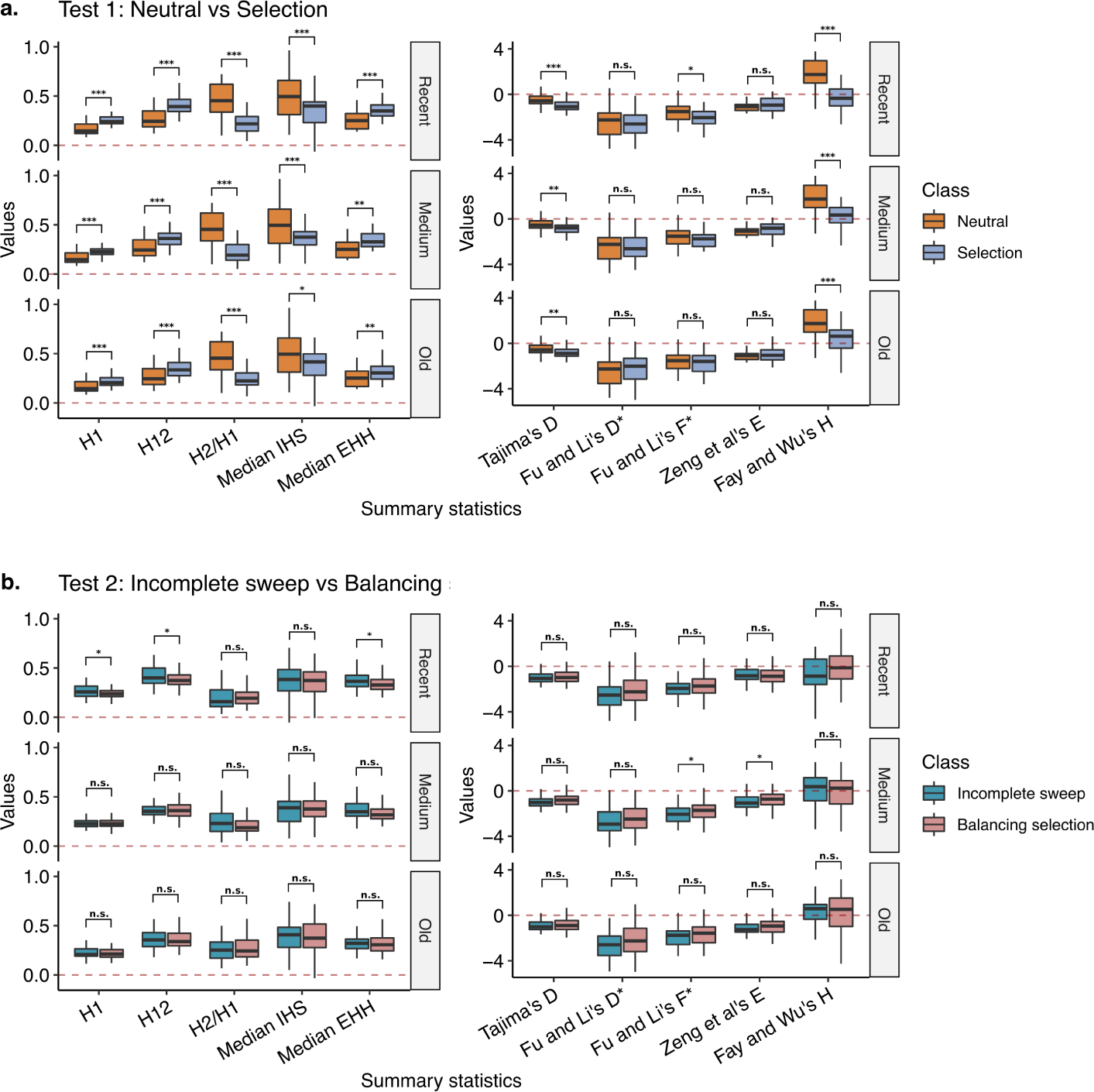
Distribution of a subset of summary statistics calculated on simulated loci under either neutral evolution or natural selection at different times of on set (recent, medium or old). Panel (a) shows the comparison between neutral evolution and natural selection (either ongoing positive selection or balancing selection). Panel (b) shows the comparison between incomplete sweep and balancing selection. Left panels group summary statistics based on haplotype diversity while right panels group summary statistics based on allele frequency. Comparisons which are statistically significant (two-sided two-sample Mann-Whitney U test) are depicted with * (*p <* 0.05), ** (*p <* 0.01), *** (*p <* 0.001), otherwise are depicted with n.s. (not significant).

Next, we tested whether summary statistics were able to distinguish between loci under incomplete sweep and balancing selection (Test 2) and, again, we compared their distributions (Figure S16). Figure 1 (lower panel b) shows the same subset of comparisons. These results suggest that only few summary statistics can discern genomic patterns created by incomplete sweep from those created by balancing selection, and only marginally. This deficiency is particularly severe for allele frequency-based summary statistics and for medium to old times of selection onset.

### 4.2 CNN has higher prediction accuracy than ANN to distinguish between incomplete sweep and balancing selection

As summary statistics do not have power to discriminate between incomplete sweep and balancing selection if considered individually, we then tested whether their predictive power increased when jointly integrated. Thus, we implemented a deep ANN which receives as input all calculated summary statistics [36] and predicts whether a given locus is under either neutrality or natural selection, either due to an incomplete sweep or balancing selection (Test 1). We compared the predictive accuracy of ANN to an approach based on convolutional layers, in form of a CNN applied to full population genomic data as an alignment of sampled haplotypes [40].

Figure 2 illustrates the performance of ANN and CNN to predict loci under different classes of evolution. The upper panel (a) on the left side shows the training loss and accuracy over epochs for classifying a locus under either neutral evolution (NE) or selection (S, Test 1). CNN showed a high loss and lower accuracy during the first few epochs, but both methods reached qualitatively similar levels of loss and accuracy after approximately ten epochs. Confusion matrices on testing data (top panel a on the right side of Figure 2) indicate similar predictive power for ANN and CNN. Recent selective events were more likely to be correctly classified than older events. For instance, we observed that the false negative rate of identifying a gene under old selection is 9% for ANN and 14% for CNN, whereas it was 4% for ANN and 1% for CNN in case of recent selection (i.e. 20k ya).

**Figure 2:**
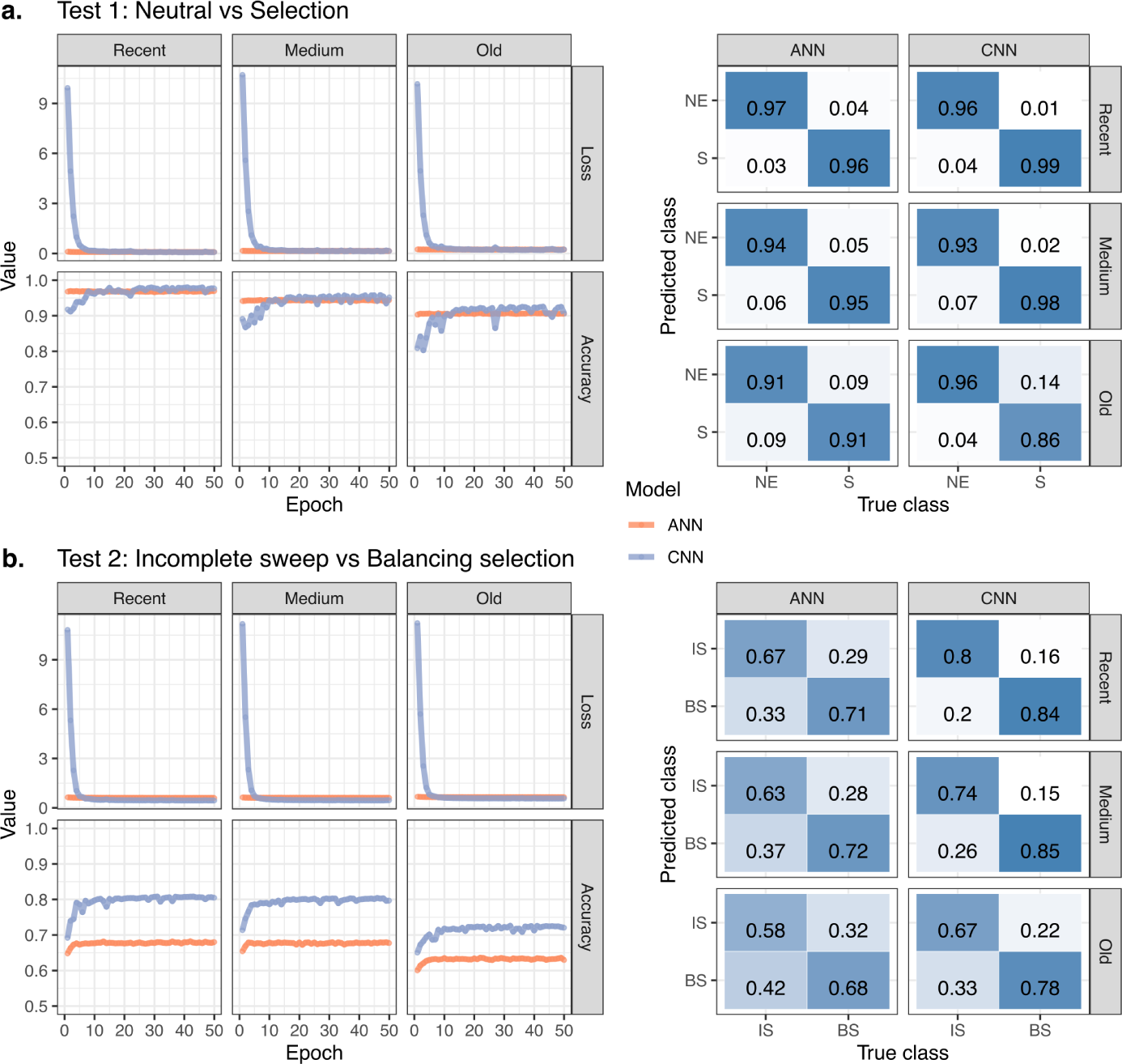
Performance of ANN (orange) and CNN (blue) to predict loci under selection (Test 1, upper panel a.) and to distinguish between incomplete sweep and balancing selection (Test 2, lower panel b.). For each category of time of onset of selection (recent, medium, old), training loss and accuracy (low-to-high gradient coloured in white-to-blue scale) over epochs are shown on the left side while confusion matrices are shown on the right side. Different classes to predict are neutrality (NE), selection (S), incomplete sweep (IS), balancing selection (BS).

The lower panel (b) of Figure 2 on the left side illustrates training loss and accuracy over epochs for classifying a locus under either incomplete sweep (IS) or balancing selection (BS, Test 2). The results recapitulated what was previously observed on the higher loss during the first few epochs for CNN. However, for this classification task, CNN exhibited a consistently higher prediction accuracy than ANN across all epochs. This observation was confirmed when investigating the confusion matrices calculated on testing data (Figure 2, right side of lower panel b). CNN consistently outperformed ANN for predicting loci under incomplete sweep or balancing selection although the overall accuracy was lower than the one obtained for Test 1. For instance, we observed a false negative rate of identifying a locus under old balancing selection of 32% for ANN and 22% for CNN, and 29% for ANN and 16% for CNN in case of recent selection. Again, recent selective events were more likely to be correctly classified than older events. However, we should stress that ANN will achieve better performance (and possibly similar prediction accuracy to the CNN) if a larger number of informative statistics are given as input. Overall, CNN had high power to identify loci under selection and substantial power to distinguish between incomplete sweep and balancing selection, two modes of evolution that leave extremely similar genomic patterns.

### 4.3 CNN is more robust than ANN to misspecified training data

The training of a neural network for population genetic inferences is conditional on a demographic and selection model to generate genomic data under different evolutionary scenarios. Therefore, we tested the robustness of both ANN and CNN to misspecified evolutionary parameters during training. Specifically, we used the already generated synthetic data and calculated the prediction accuracy for identifying loci under selection (Test 1) and for distinguishing between incomplete sweep and balancing selection (Test 2) when both ANN and CNN were trained on a specific time of onset of selection (recent, medium, old) but tested on a different value. By doing so, we were able to quantify any drop in accuracy when the training data did not reflect the underlying true evolutionary model.

Figure 3 shows the prediction accuracy for both tests (Test 1 and Test 2, on columns) and networks (ANN and CNN, on rows) for all possible pairs of time of onset of selection between training and testing data. Numbers on the antidiagonal represent accuracy values when the model used for both training and testing was the same. Numbers outside the antidiagonal indicate accuracy values when the models employed for training and testing differed. We observed a marginal decline in accuracy when using incorrect training data for Test 1 for both networks which performed similarly. These results were confirmed when investigating all corresponding confusion matrices (Figure S17). For Test 2, the drop in accuracy when employing a different model for training was more evident than for Test 1, although CNN outperformed ANN in most scenarios (Figures 3, S18).

**Figure 3:**
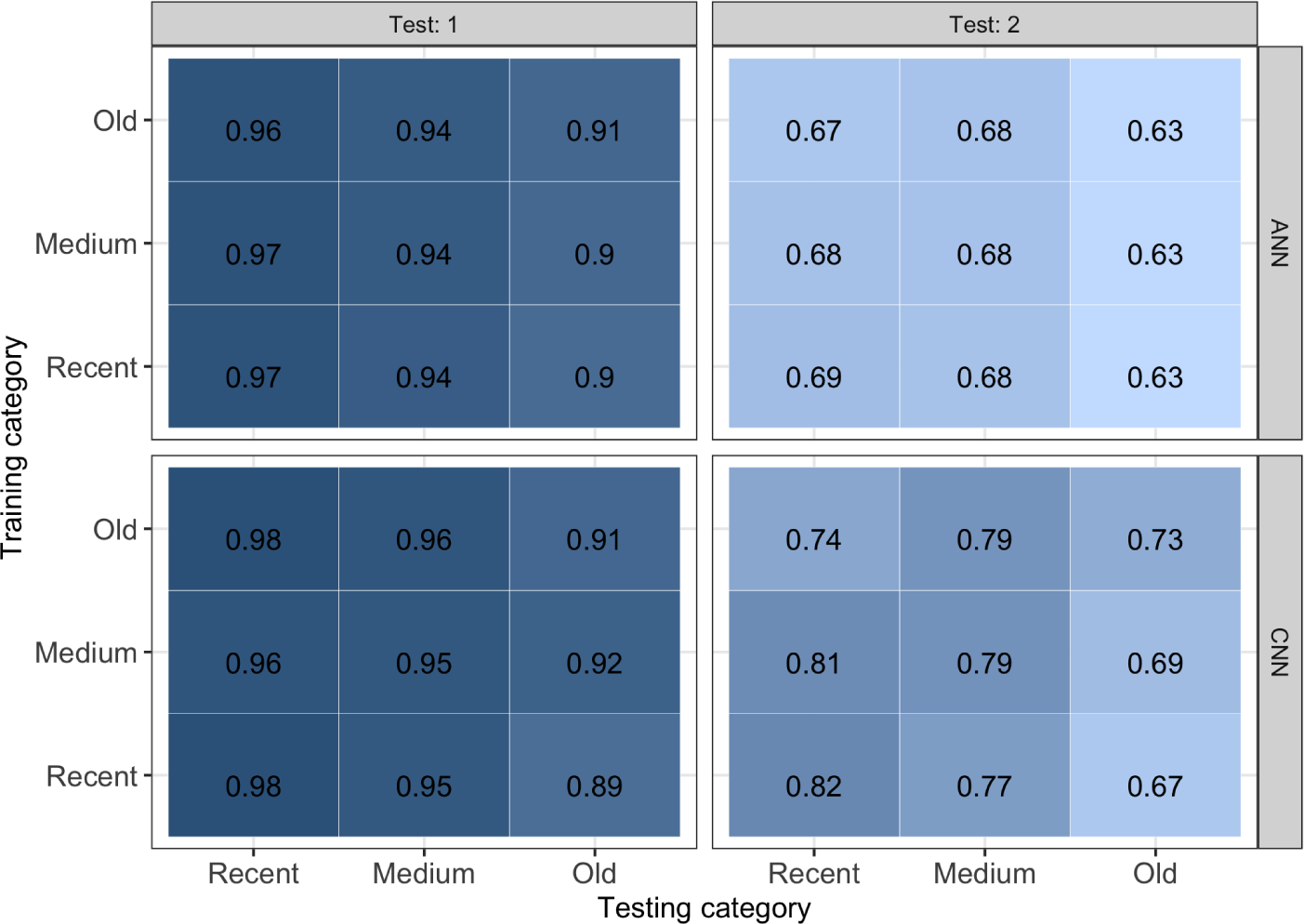
Prediction accuracy (low-to-high gradient coloured in white-to-blue scale) for classifying loci under different evolutionary events (Test 1 and Test 2, on columns) and methods (ANN and CNN, on rows) for all pairs of classes for time of onset of selection between training (y-axis) and testing data (x-axis). The antidiagonal shows accuracy values when the model used for both training and testing is the same, while accuracy values outside the antidiagonal are obtained when the models employed for training and testing differ.

To further test the robustness of our inferences to a misspecified model, we tested both architectures trained on a European population to simulated data generated from a demographic model of African and East Asian populations [48]. Accuracy values for Test 1 are marginally affected by a misspecified demographic model (Figures S19, S20), while we observed a slightly more pronounced decrease in performance for Test 2 (Figures S19, S20).

### 4.4 CNN identifies signatures of recent natural selection in *MEFV* gene

We deployed the trained networks, both ANN and CNN, on genomic data for the *MEFV* gene from CEU population from the 1000 Genomes Project [49]. MEFV gene has been previously associated with both balancing selection [17] and ongoing positive selection [46]. Here we tested whether MEFV gene has been targeted by natural selection and, if so, whether by balancing selection or incomplete sweep.

To assess the false positive rate, we extracted flanking genomic regions to *MEFV* predicted to be under neutral evolution [70], and deployed both ANN and CNN algorithms on all intermediate frequency variants. We expected the networks not to predict signals of selection within these control neutral regions. ANN predicted 23 out of 42 sites to be under selection regardless of the time of onset of selection (Figure S21). Therefore, we decided not to use the ANN algorithm for inferences on the *MEFV* gene, as it showed a high false positive rate based when applied to putative neutral genomic regions. In contrast, CNN provided strong support for 39 out of 42 sites to be under neutral evolution, with only three sites possibly predicted to be under selection regardless of the time of onset (Figure S22).

Next, we aimed to identify signals of natural selection and deployed the trained CNN within the *MEFV* genomic region of European samples (CEU) from the 1000 Genomes Project database [49]. We observed a large proportion of sites with intermediate allele frequency predicted to be under natural selection (Test 1) regardless of the time of onset of selection (Figure 4, upper panel). All sites under selection were predicted to be under incomplete sweep rather than balancing selection (Figure 4, second panel from top).

**Figure 4:**
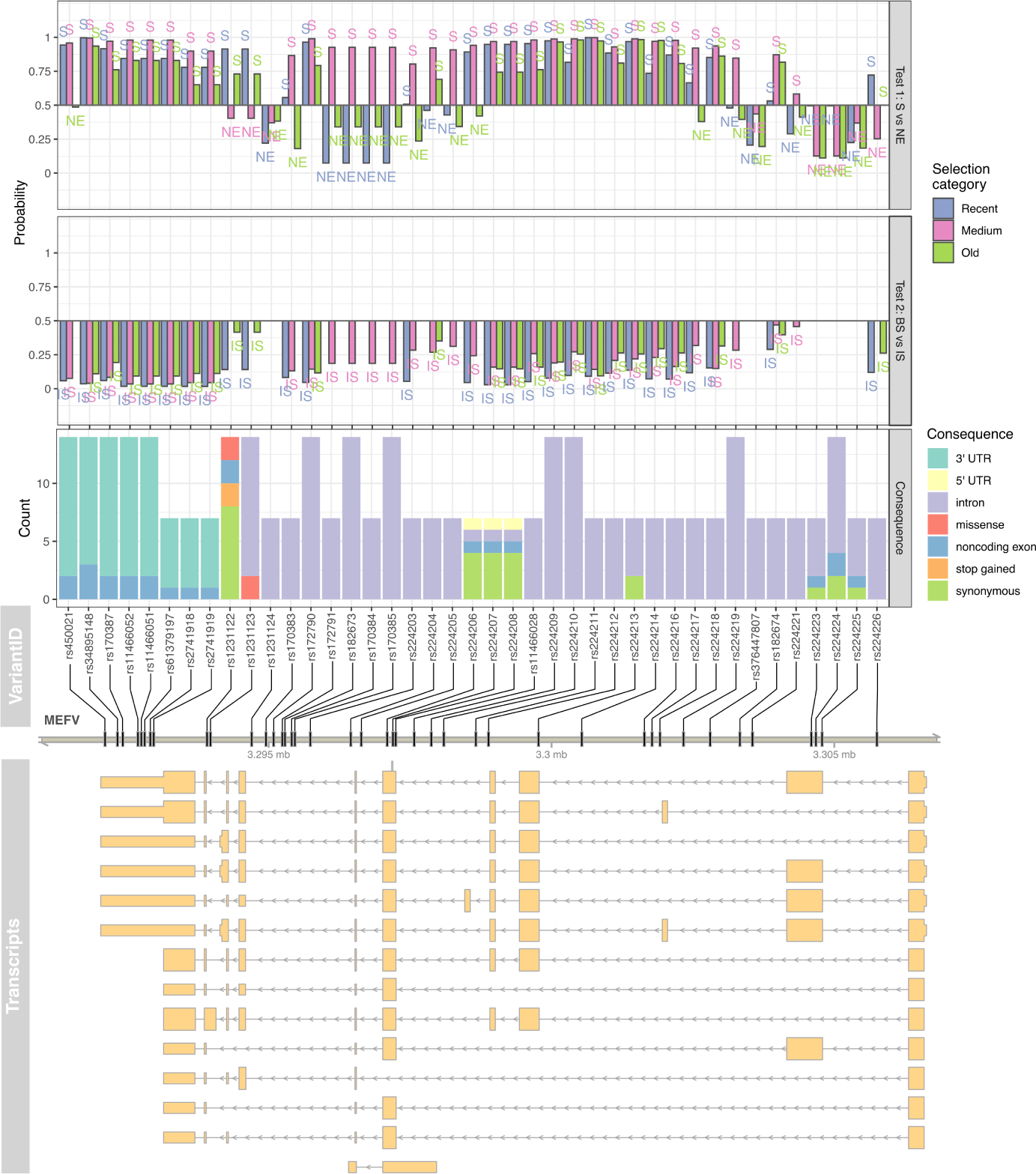
Prediction of sites under natural selection (Test 1, upper panel) or balancing selection *vs.* incomplete sweep (Test 2, second panel from top) on intermediate-frequency variants in the *MEFV* gene for a European population. For each tested variant, the predicted functional impact across all isoforms is reported (counts of functional consequences on third panel, genomic location on fourth panel and transcripts on fifth panel from the top).

Sites predicted to be under selection (or in LD with the target of selection) encompass a haplotype block spanning from intron 2 to 3’ UTR (untranslated region, Figure S23). Most of these variants are possibly functionally silent as they lay within introns or represent synonymous substitutions (Figure 4, third to fifth panels from top). However, two mutations within this region represent either missense (rs1231123, rs1231122) or stop-gained (rs1231122) substitutions, depending on the corresponding isoform. The predicted signals of selection in the *MEFV* gene were confirmed when deploying the trained network to genomic data from TSI samples [49], another European population (Figure S24). However, the results obtained using TSI population showed a higher false positive rate when deployed to neutral genomic regions (Figure S25) than the ones obtained using CEU population, possibly because the network was trained on simulated data conditional on a demographic model inferred for the CEU population. In fact, 7, 14 and 10 out of 38 neutral sites were predicted to be under selection with recent, medium and old time of onset, respectively, using TSI population. In contrast, 3, 13 and 9 out of 42 neutral sites were labelled as targets of selection with recent, medium and old time of onset, respectively, using CEU population.

## 5 Discussion

In this study we demonstrated the utility of deep learning to identify genomic signals of recent natural selection on intermediate frequency variants. We showed that algorithms based on either summary statistics (i.e. ANN) or full genomic data (i.e. CNN) had comparably high power to infer selective regimes (Figure 2) and exhibit lower false positive and false negative rates than commonly used neutrality tests (Figure S26). However, CNN had higher accuracy to distinguish between loci under balancing selection and incomplete sweep (Figure 2), it was generally more robust to incorrect training data (Figure 3), and it had a lower false positive rate when deployed on neutral genomic regions than ANN (Figures S21-S22). Finally, we illustrated the applicability of deep neural networks to detect and characterise signals of natural selection on common variants within the *MEFV* gene region (Figure 4).

Our results on the high predictive power offered by deep learning, and specifically by convolutional neural networks, to detect signals of natural selection expand previous findings [39, 38, 40, 41] to cases where the beneficial allele is at intermediate frequency. CNN outperformed ANN to distinguish between incomplete sweep and balancing selection although, in our analyses, its training was slower by a factor of 300. In fact, CNN had more than 4 million parameters to estimate, in contrast to ANN which had approximately 2,000. Additionally, ANN received as input informative features (i.e. summary statistics) while convolutional filters in the CNN learned the optimal features from the raw data whilst training. In machine learning, the design of such features had been a major part of information engineering. As an illustration, in the field of computer vision, the ”features” used for many practical algorithms until the early 2000s consisted of hand-engineered gradient estimators [74], typically at multiple spatial scales [75, 76], applied to images (arrays of pixels). The observation that features emerge within a deep network has been repeated in different domains. Therefore, we envisage that a novel area of research will focus on extracting informative features from trained networks for population genetic inference, possibly by analysing activation or saliency maps [77]. It is important to note that ANN will achieve higher performance with the inclusion of additional summary statistics not considered herein.

This study also contributes to ongoing efforts to design architecture and devise training techniques for deep learning algorithms in population genetics [41]. Resizing images to smaller dimensions appeared to reduce overfitting and learning time (Figure S9) and could be considered a complementary strategy to approaches based on cropping or padding [38]. The strategy to separately sort rows based on the presence or absence of the putative target variant is an alternative solution to adopt more general, but computational expensive, architectures based on exchangeable neural networks [39, 41]. We also explored the applicability of forward-in-time simulations to train deep neural networks for population genetics and the usefulness of data augmentation (Figure S10) to reduce the computational time required to generate synthetic training data. The use of forward-in-time simulations should generate more realistic synthetic population genomic data and model more complex evolutionary scenarios than by using coalescent simulations. In any case, as suggested in this study (Figures S21-S22), false positive and negative rates should be assessed by deploying trained networks on loci previously identified as targets of selection or neutrally evolving. Further research on the design of neural networks for population genetics is in need, for instance by maximising the prediction accuracy over a grid of hyperparameters (e.g., number of units and layers) using Neural Architecture Search [78].

We show that deep neutral networks achieved higher prediction power to differentiate between the effects of neutral evolution, balancing selection and incomplete sweep for variants segregating at intermediate frequency (Figure 2) than commonly used summary statistics (Figure 1). However, the accuracy to distinguish between incomplete sweep and balancing selection using CNN ranges from 72% to 80% depending on the time of onset of the selection, with more recent events (around 20k ya) more accurately classified (Figure 3). While this accuracy is far higher than that achieved using summary statistics, higher accuracy could be achieved by employing a larger training data set, by using more extensive hyper-parameter tuning and architecture search, and by treating overdominance and negative frequency dependent selection as separate prediction categories. In fact, future extensions of this study will include testing to distinguish between overdominance and negative frequency-dependent selection once a variant is predicted to be under balancing selection. It is likely that a different CNN architecture and training data is needed for this purpose as, for instance, information on heterozygosity (not considered herein given the simulation strategy) will likely emerge as an important feature. Additionally, a wider spectrum of times of the onset of selection should be considered to assess the power to predict balancing selection at different evolutionary scenarios. Finally, the CNN herein proposed requires fully resolved haplotypes with no missing data. Furthermore, we argue that such approach should be locus-specific as the network needs to be trained with the local characteristics of the region of interest (e.g., recombination rate). Therefore, this CNN is more suitable to be deployed to deep resequencing data on single loci of interest rather than to genome-wide low-coverage sequencing data. In the latter case, we argue that an ANN receiving as input statistical estimates of summary statistics from genotype likelihoods [79] might be a valuable alternative. Nevertheless, the effect of data uncertainty should be further explored.

The analyses on the *MEFV* gene performed herein complement previous findings [80] to suggest that this gene has been subjected to different evolutionary forces. The *MEFV* gene encodes for the Pyrin protein which plays an important role in inflammatory processes [81]. Five different functional domains have been identified within the Pyrin protein. The PYD domain (aa 1-92) is present in at least 20 human proteins involved in inflammatory pathways. However, in the analyses we performed the PYD domain seems to have neutrally evolved. The Pyrin central region hosts three domains: a bZIP domain (aa 266-280), a B-box domain (aa 370-412) and a coiled-coil domain (CC, aa 420-440). The role of these three domains has not been thoroughly elucidated and few FMF-causing variants localize to Pyrin’s central region [82, 83]. Nevertheless, from our data this central region is apparently under recent selection (Figure 4) or is in LD with beneficial alleles (Figure S23). Similarly, the B30.2 domain (also known as PRY/SPRY domain), which is encoded by the *MEFV* exon 10 where most of the FMF-causing variants cluster [84] shows the same genetic patterns of ongoing selection.

A recent study demonstrated that the FMF-associated variants M694V, M680I and V726A, all localizing to the B30.2 region, decrease the binding of *Yersinia pestis* virulence factor YopM [46]. Further, the authors provided evidence that M694V and V726A variants were subject to recent positive selection in a cohort of Turkish individuals. Finally, FMF knock-in mice demonstrated survival advantage compared to wild type mice. Thus, these experimental evidences suggest that mutations in the human Pyrin may have conferred resistance to *Yersinia pestis* [46]. However, the possibility that other pathogens could have concurred in conferring a selective advantage cannot be ruled out. Indeed, contrary to previous claims of overdominance acting on *MEFV* [17], our new results and Park *et al.*’s study suggest that the selection on human Pyrin is directional and either recent or possibly still ongoing. In fact, the frequency of M694V and V726A kept rising [46] although no plague outbreaks rose to the scale of a pandemic after the 17*^th^* century.

The population sample we analysed in this study is different from the Turkish cohort investigated by Park *et al.* which overlaps significantly with one of the plague outbreak sites. Nevertheless, even in the different population sample we analysed, the data presented herein suggest signals of recent selection on the human Pyrin. While our computational predictions are unable to identify the causal variant, it is possible to hypothesise that Pyrin, specifically its B30.2 region, could confer resistance to a broader range of pathogens including those causing more recent pandemics. Likewise, we cannot rule out more complex evolutionary scenarios, with *MEFV* being subjected to long-term balancing selection and recent positive selection on standing variation. A comprehensive picture of ongoing selection signatures in *MEFV* could be achieved by deploying deep neural networks trained on variants segregating at low or high frequency and to a wide range of Mediterranean populations. Finally, additional power to characterise recent selection in *MEFV* could be gained by integrating data from ancient genomes [85] as this would be particularly suitable to relate adaptation to past epidemics to current pathogenic threats [86].

In this study we demonstrated how deep learning, and in particular convolutional neural networks, were able to perform predictions currently inaccessible by commonly used strategies based on summary statistics. In particular, we showed that deep neutral networks can differentiate between signals of incomplete sweep and balancing selection, despite the two evolutionary events leaving qualitatively similar patterns of genetic variation. Furthermore, our application to detect signals of selection on FMF-associated alleles highlighted the importance of a population genetic approach to understand the molecular basis of susceptibility and/or resistance to infectious diseases.

## 6 Acknowledgements

This work was supported by a Leverhulme Trust Research Grant (RPG-2018-208) and an Imperial College European Partners Fund to MF. We acknowledge the support offered by the Erasmus+ programme to UI. We are grateful to Aida Andŕes, Anil A. Bharath and Mehmet Somel for discussions and three anonymous reviewers for comments on the manuscript. We also thank Kivilcim Basak Vural and the METU Comparative and Evolutionary Biology Group for computational support.

## 8 Data Accessibility

Detailed tutorials on pipelines for training and prediction, along with all the scripts used in this study, are available within *BaSe* package at https:// github.com/ulasisik/balancing-selection. Sequencing data on human populations were retrieved from The International Genome Sample Resource (IGSR) at https://www.internationalgenome.org.

## 9 Author Contributions

MF and UI designed the research. UI performed the research with contributions from AS. MF, UI and AS analyzed data and wrote the paper.

## 10 Abbreviations

ANN: artificial neural network bp: base pairs
BS: balancing selection
CNN: convolutional neural network
IS: incomplete sweep
LD: linkage disequilibrium
ML: machine learning
NE: neutral evolution
ReLU: Rectified Linear Units
S: natural selection
TSI: Tuscan in Italy
UTR: untranslated region
ya: years ago

## 11 Tables and Figures (with captions)

## 12 Supplementary Tables and Figures (with caption)

Table S1: Parameters used in the demographic model to simulate genomic data from Table 2 in Jouganous *et al.* [48]. Parameters are defined in Gravel *et al.* [87].

Table S2: Parameters to simulate intermediate frequency alleles under different tested scenarios of selection. For each case, we report the used values of time of onset, selection coefficient, dominance and coefficient and parameters for relative fitness (A and B, as *f itness* = *A − B × f requency*).

Table S3: Summary statistics calculated as input to the ANN. Each statistic is calculated on two regions surrounding the target variant.

Table S4: Parameters of the optimised CNN architecture. Layer notations: I=Input, C=Convolution, BN=Batch Normalization, P=Pooling, A=Activation(ReLU), D=Droupout, F=Flatten, FC=Fully-Connected(Dense), O=Output.

**Figure S1:**
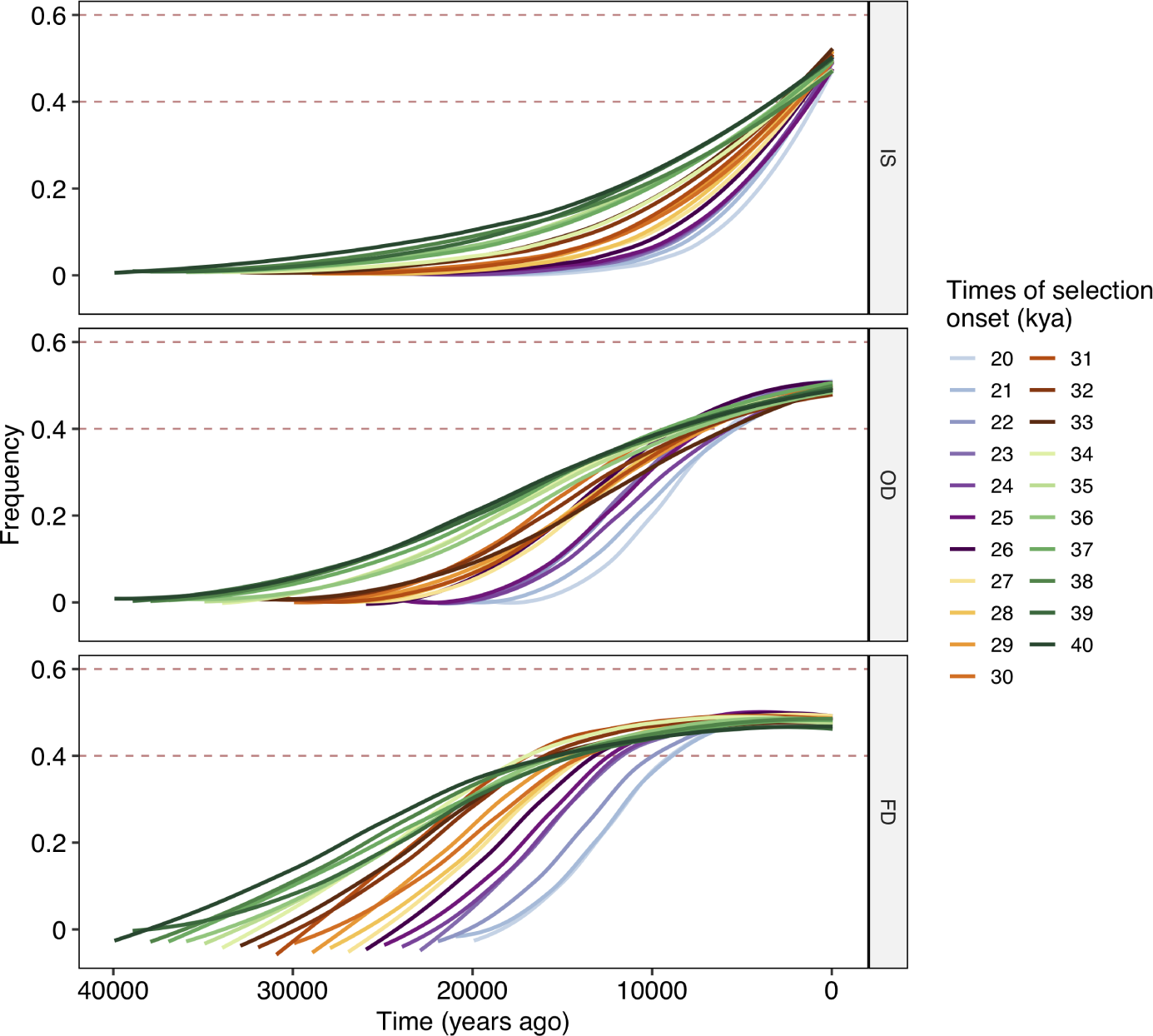
Examples of simulated allele frequency trajectories for different times of onset and different modes of selection: incomplete sweep (IS), overdominance (OD), negative frequency dependent selection (FD).

**Figure S2:**
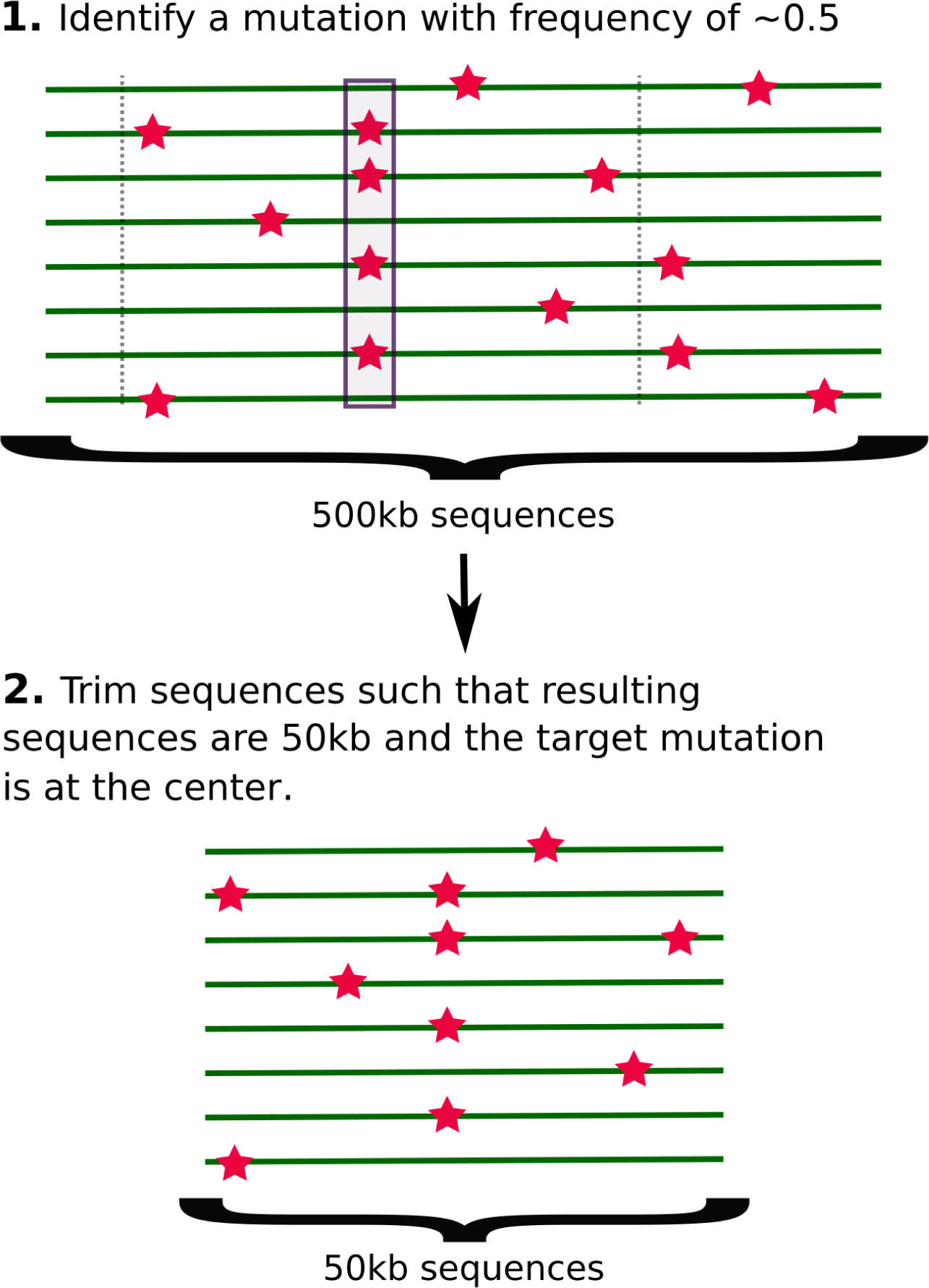
A cartoon illustrating the strategy to generate simulations of neutral regions with intermediate frequency alleles.

**Figure S3:**
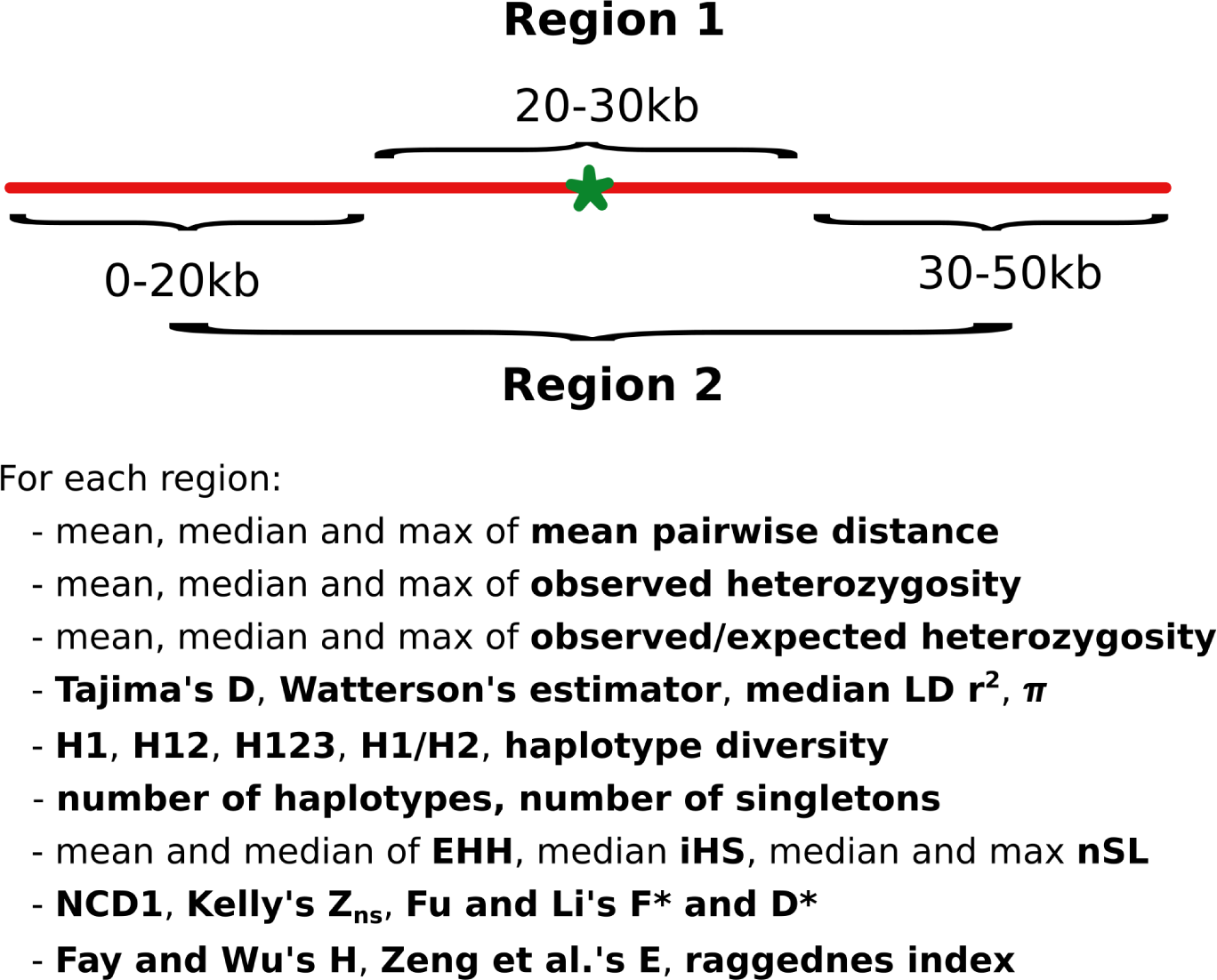
A cartoon illustrating the strategy to calculate all summary statistics used. For each 50k bp locus, each statistic is calculated on two regions, a proximal one labelled 1 (20-30k bp) and a distal one labelled 2 (0-20k bp + 30-50k bp).

**Figure S4:**
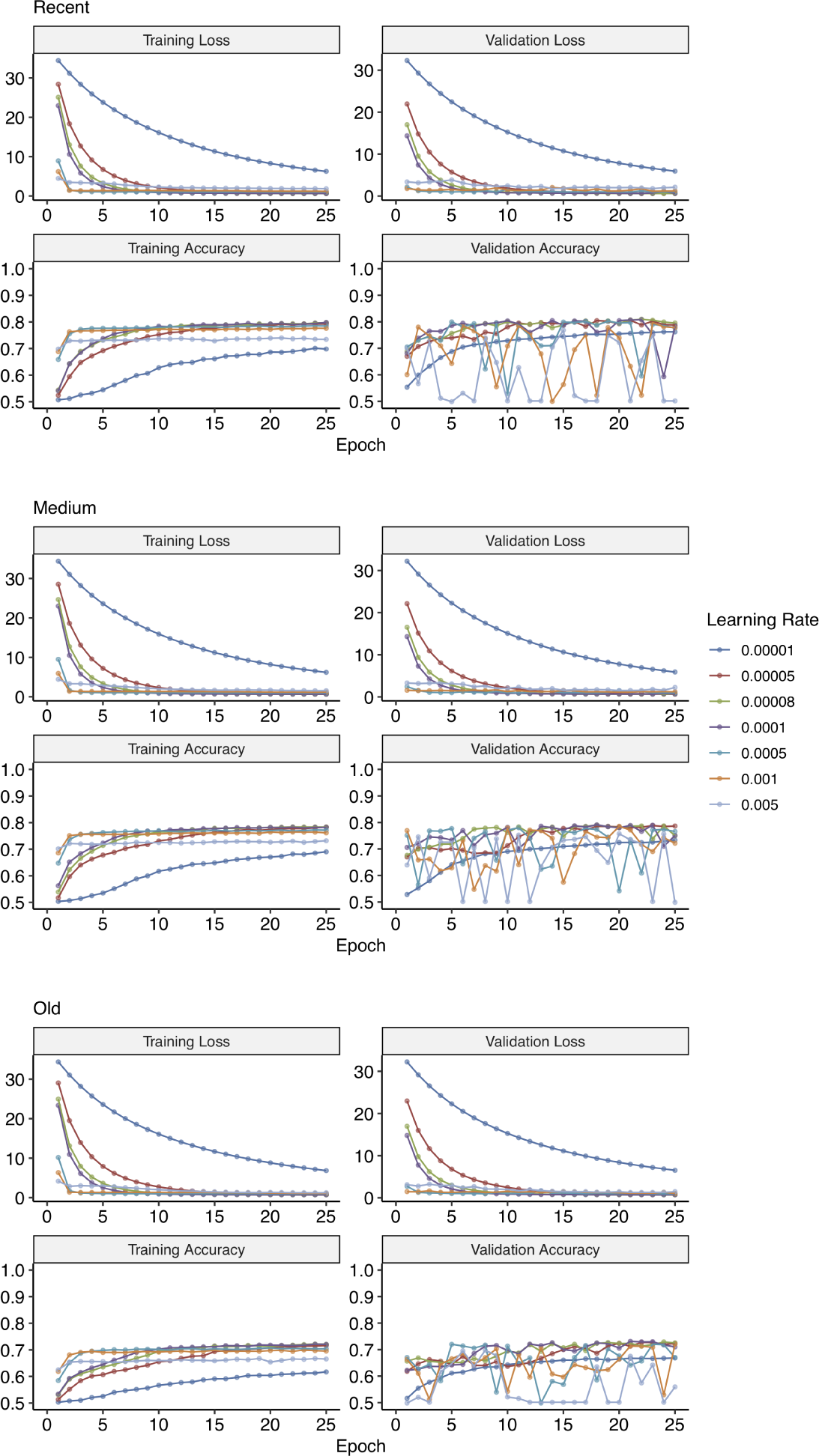
Training and validation loss and accuracy plots for hyper-parameter tuning of learning rate to train CNN for Test 2 (incomplete sweep *vs.* balancing selection) at different times of onset of selection (see Methods).

**Figure S5:**
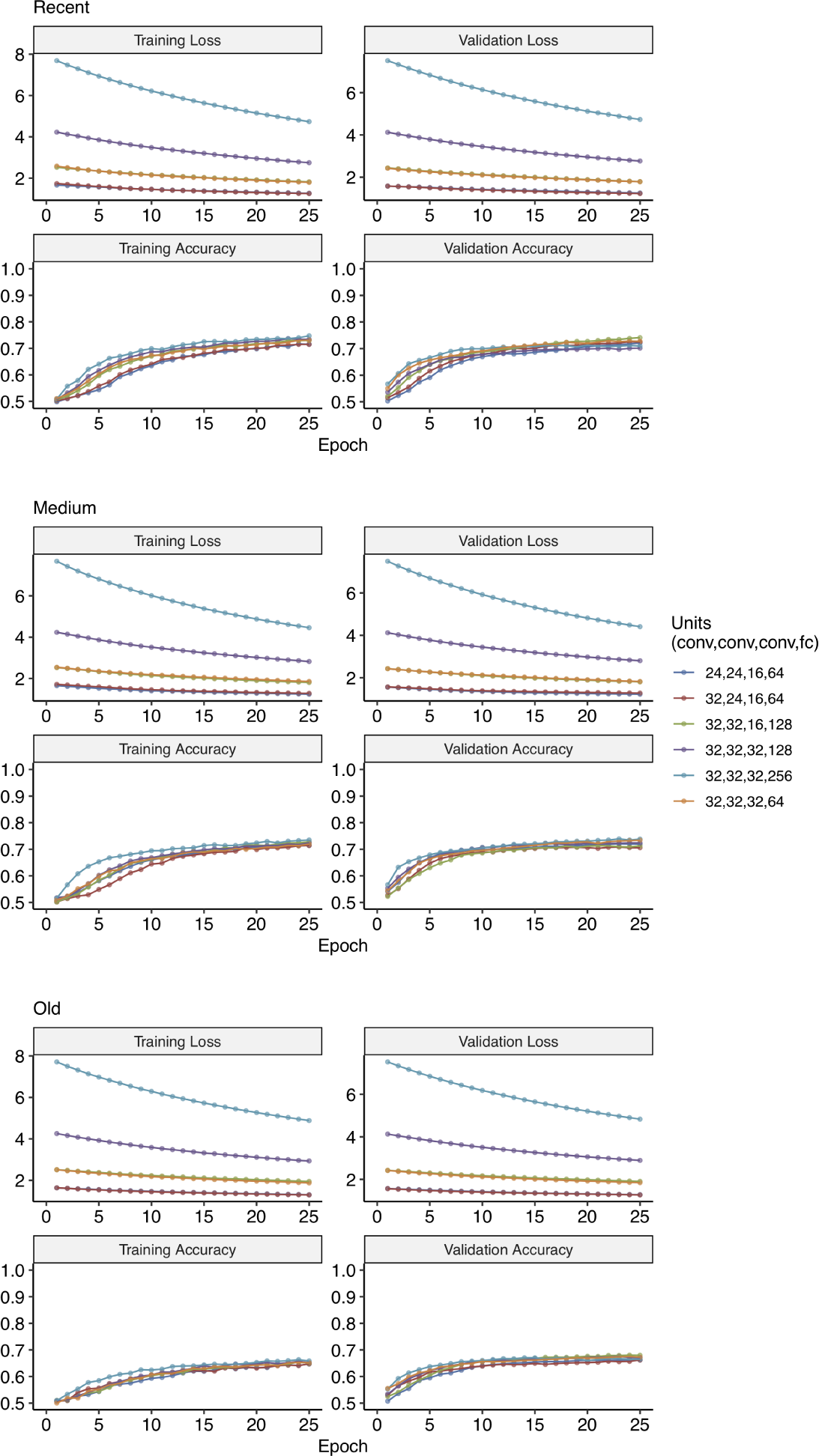
Training and validation loss and accuracy plots for hyper-parameter tuning of number of units for convolutional (conv) and fully-connected (fc) layers to train CNN for Test 2 (incomplete sweep *vs.* balancing selection) at different times of onset of selection (see Methods).

**Figure S6:**
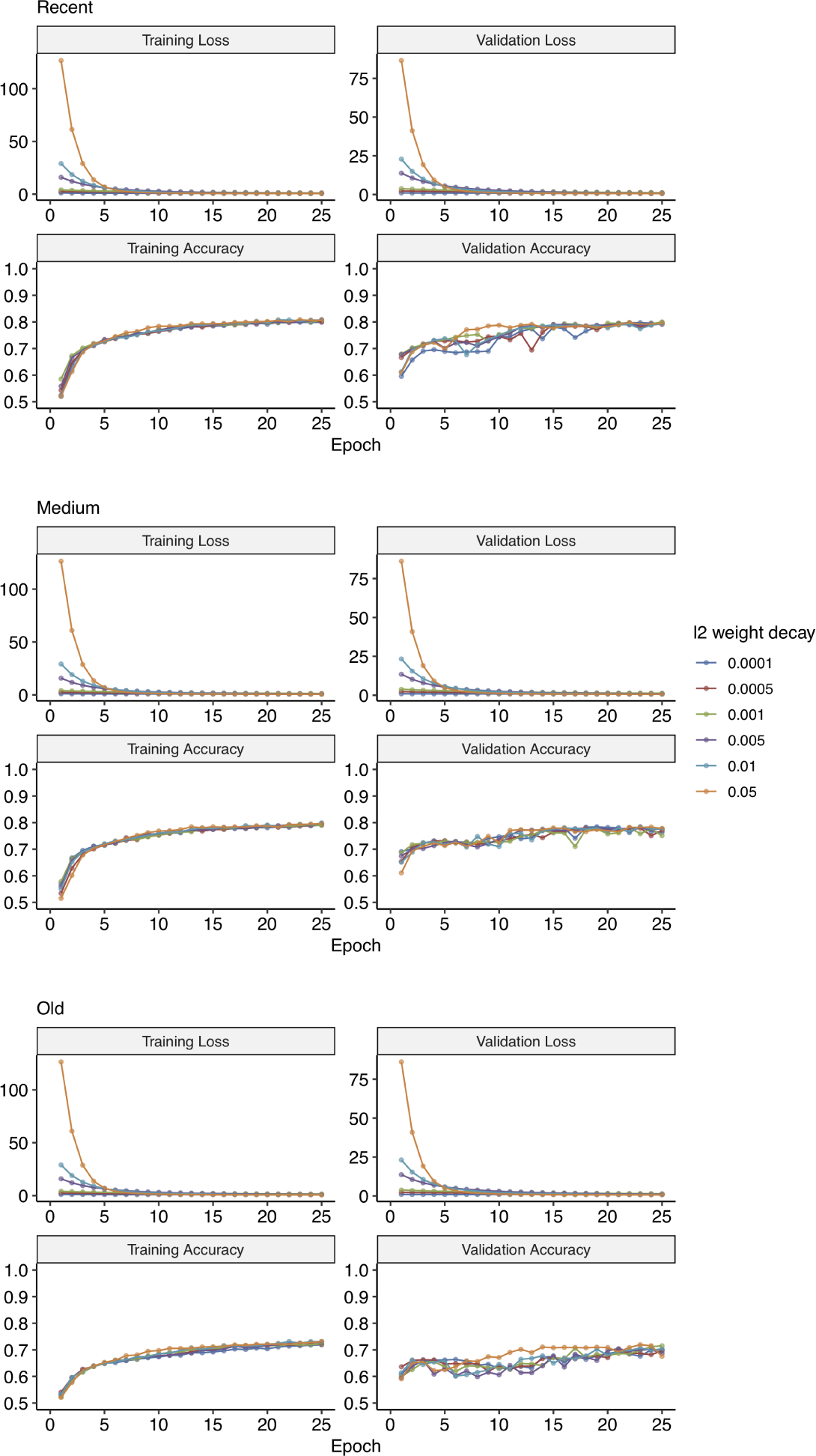
Training and validation loss and accuracy plots for hyper-parameter tuning of regularisation rates to train CNN for Test 2 (incomplete sweep *vs.* balancing selection) at different times of onset of selection (see Methods).

**Figure S7:**
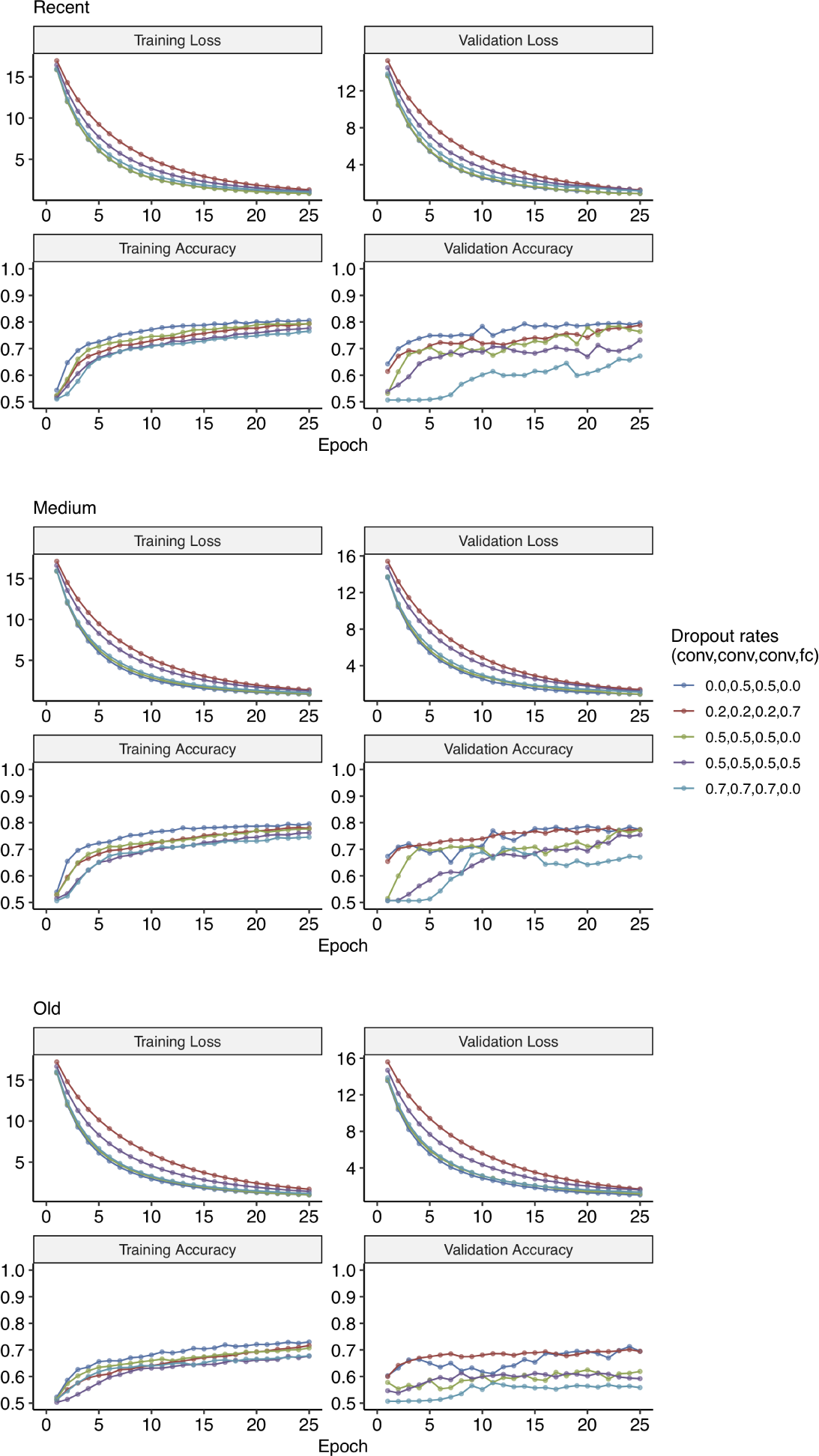
Training and validation loss and accuracy plots for hyper-parameter tuning of dropout rates for convolutional (conv) and fully-connected (fc) layers to train CNN for Test 2 (incomplete sweep *vs.* balancing selection) at different times of onset of selection (see Methods).

**Figure S8:**
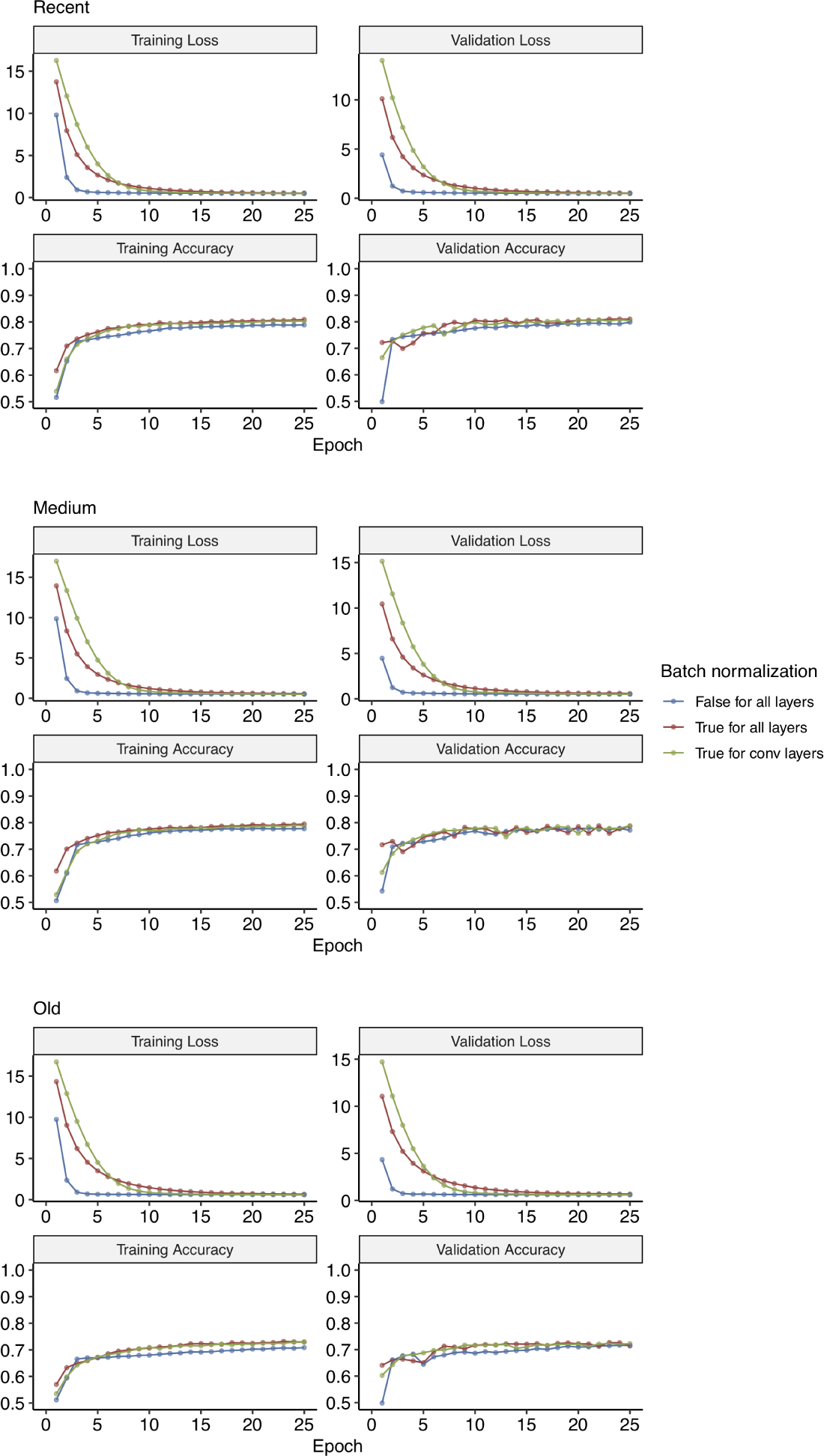
Training and validation loss and accuracy plots for hyper-parameter tuning of batch normalisation to train CNN for Test 2 (incomplete sweep *vs.* balancing selection) at different times of onset of selection (see Methods).

**Figure S9:**
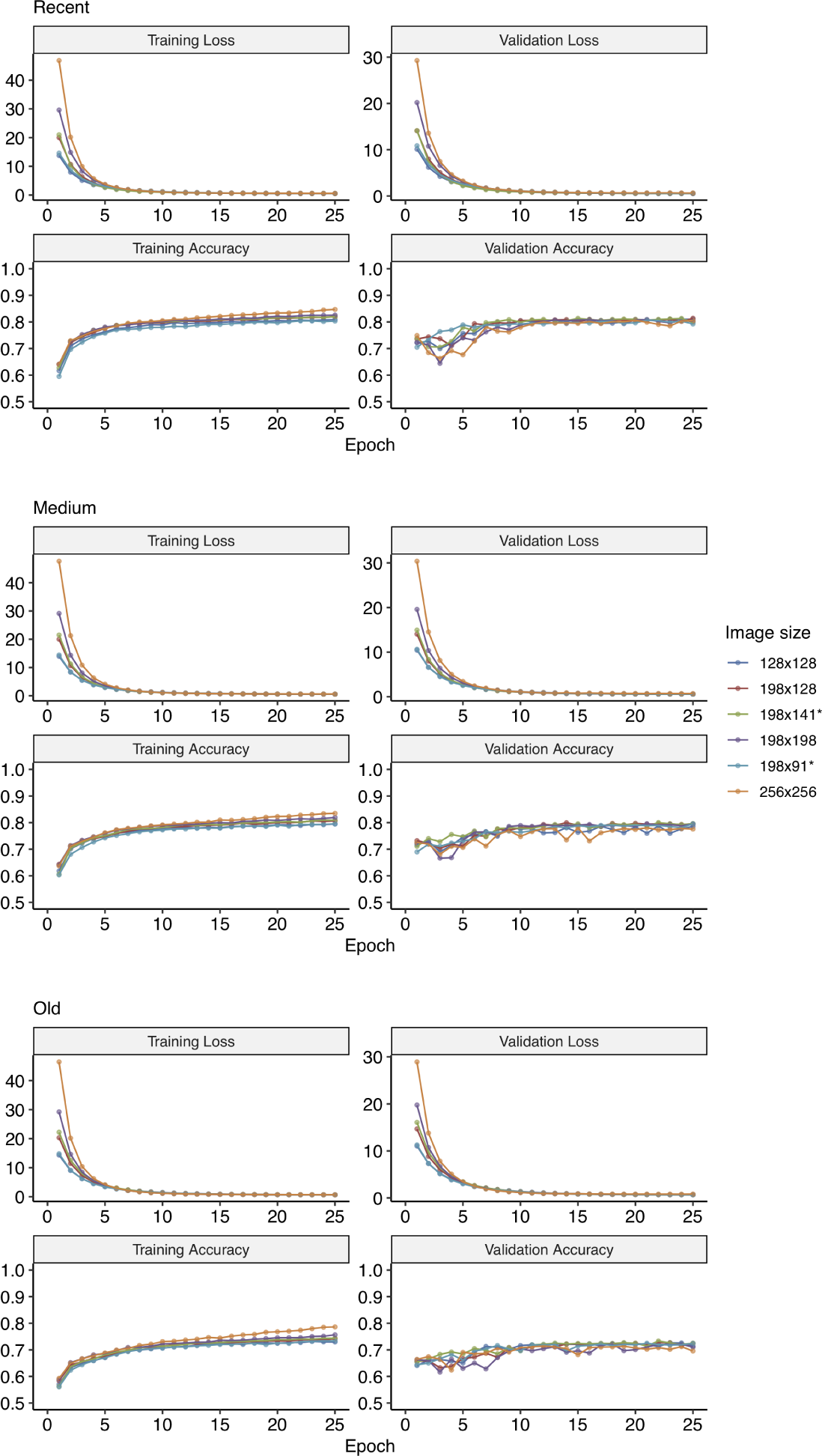
Training and validation loss and accuracy plots for hyper-parameter tuning of reshaping images to train CNN for Test 2 (incomplete sweep *vs.* balancing selection) at different times of onset of selection (see Methods).

**Figure S10:**
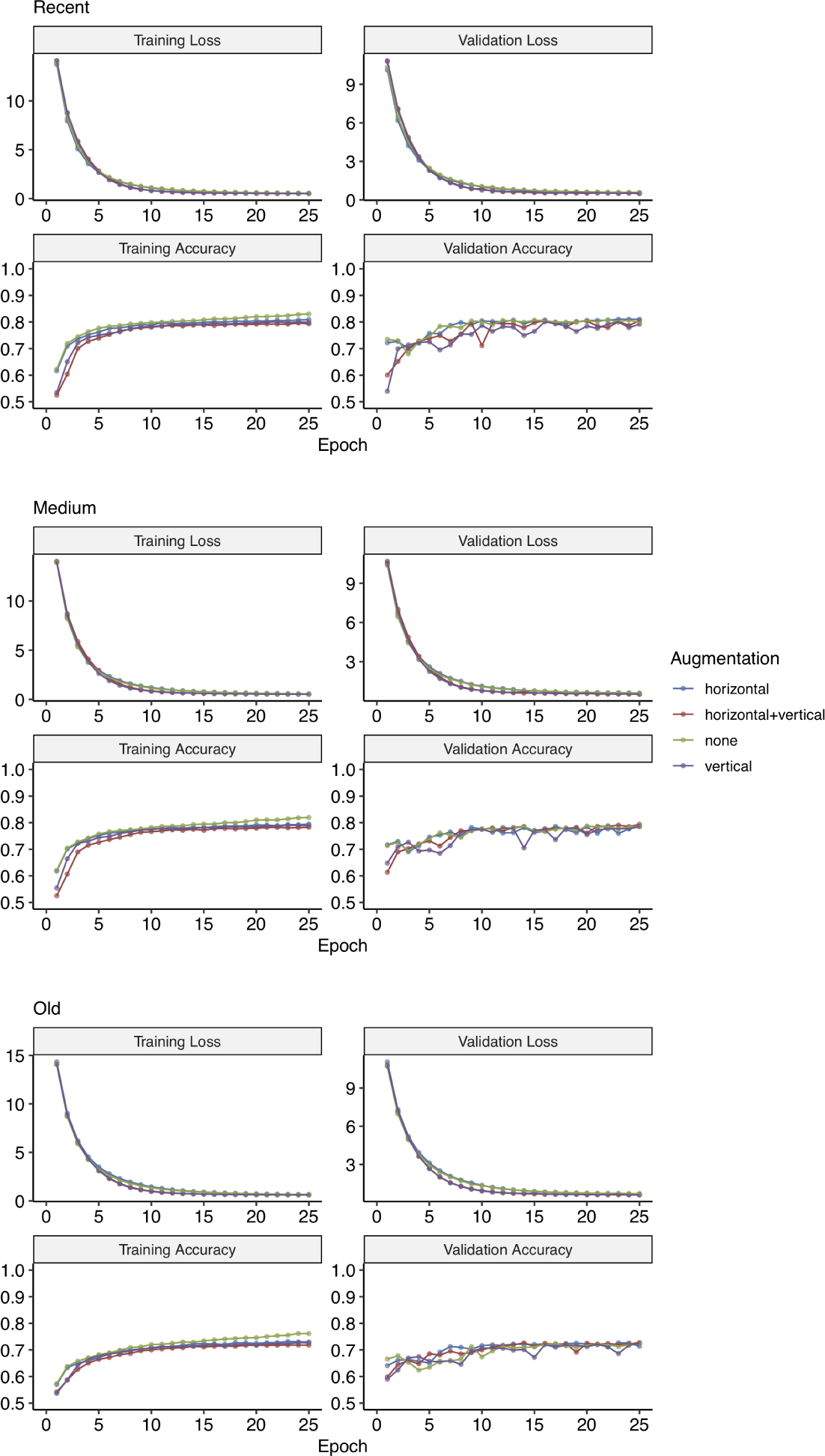
Training and validation loss and accuracy plots for hyper-parameter tuning of data augmentation (i.e. flipping images) to train CNN for Test 2 (incomplete sweep *vs.* balancing selection) at different times of onset of selection (see Methods).

**Figure S11:**
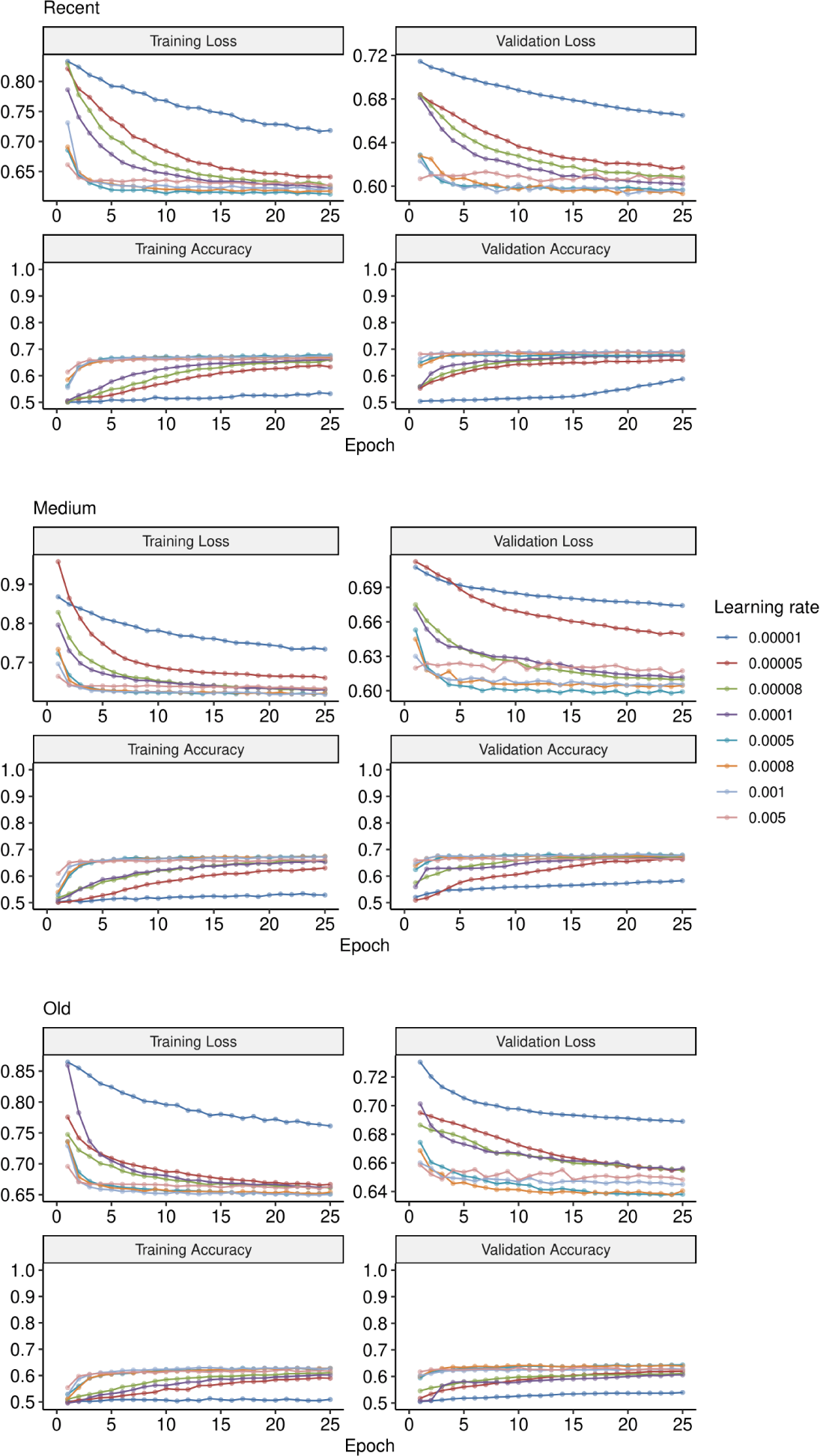
Training and validation loss and accuracy plots for hyper-parameter tuning of learning rate to train ANN for Test 2 (incomplete sweep *vs.* balancing selection) at different times of onset of selection (see Methods).

**Figure S12:**
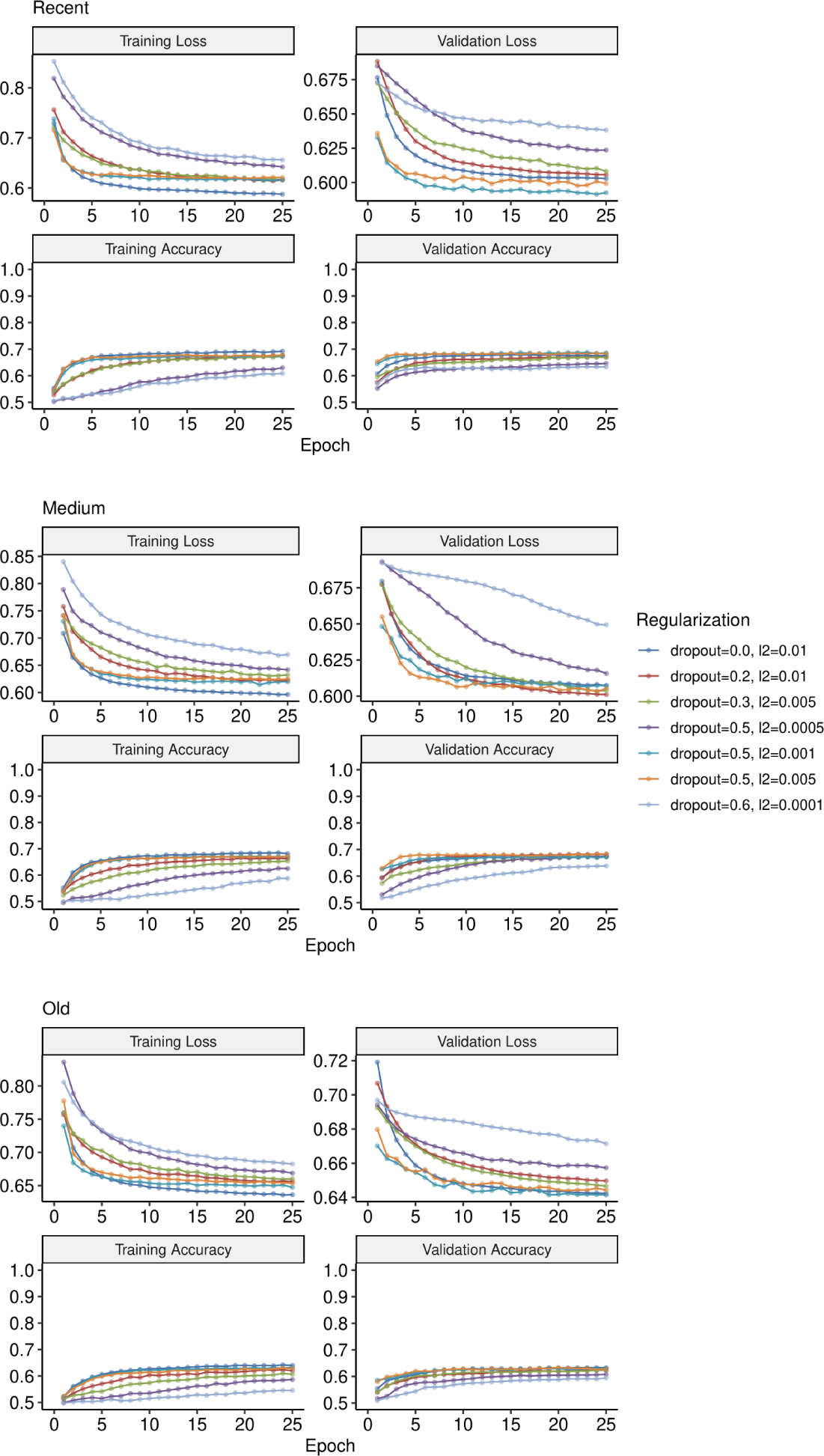
Training and validation loss and accuracy plots for hyper-parameter tuning of regularisation rates to train ANN for Test 2 (incomplete sweep *vs.* balancing selection) at different times of onset of selection (see Methods).

**Figure S13:**
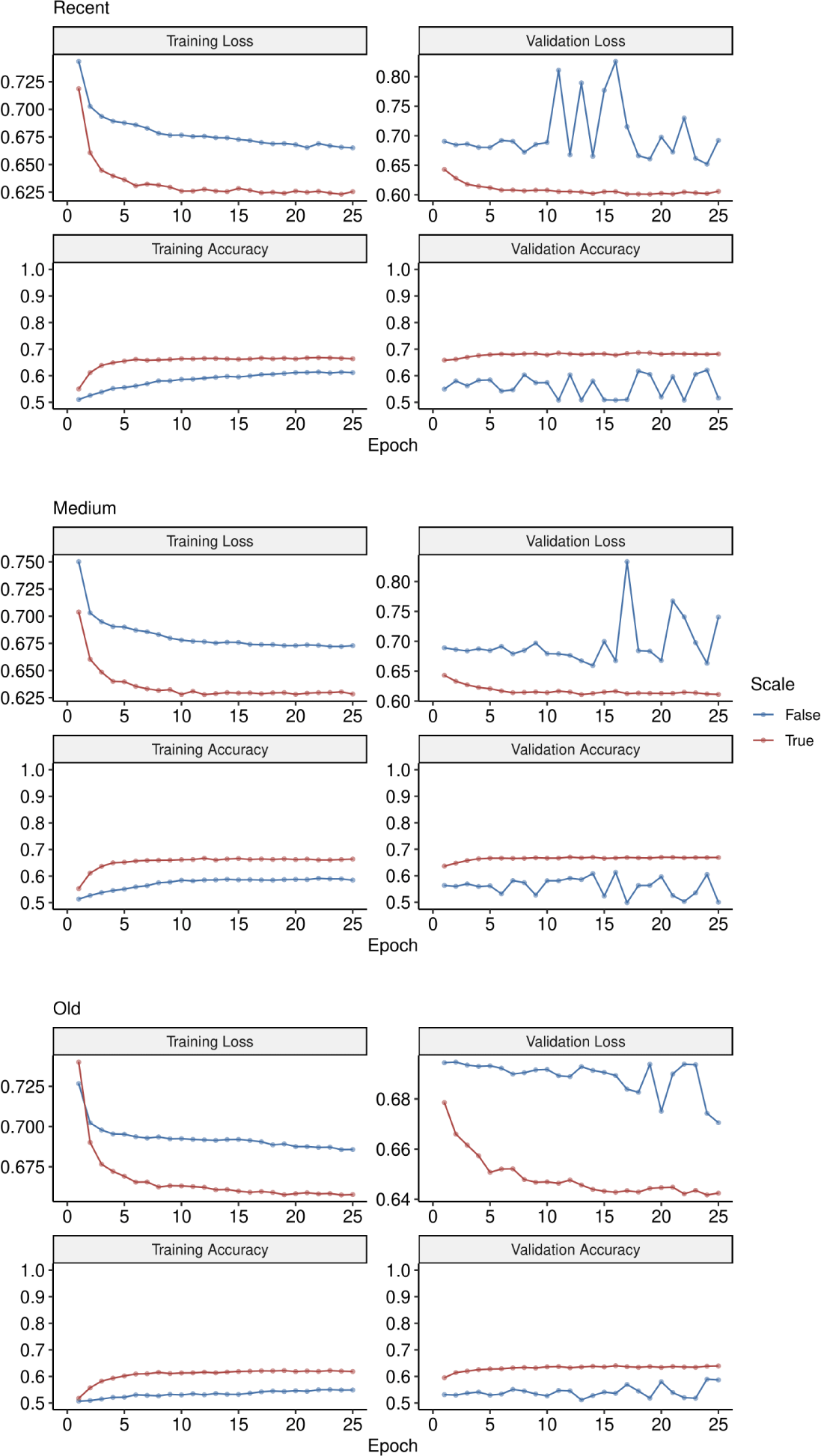
Training and validation loss and accuracy plots for hyper-parameter tuning of scaling to train ANN for Test 2 (incomplete sweep *vs.* balancing selection) at different times of onset of selection (see Methods).

**Figure S14:**
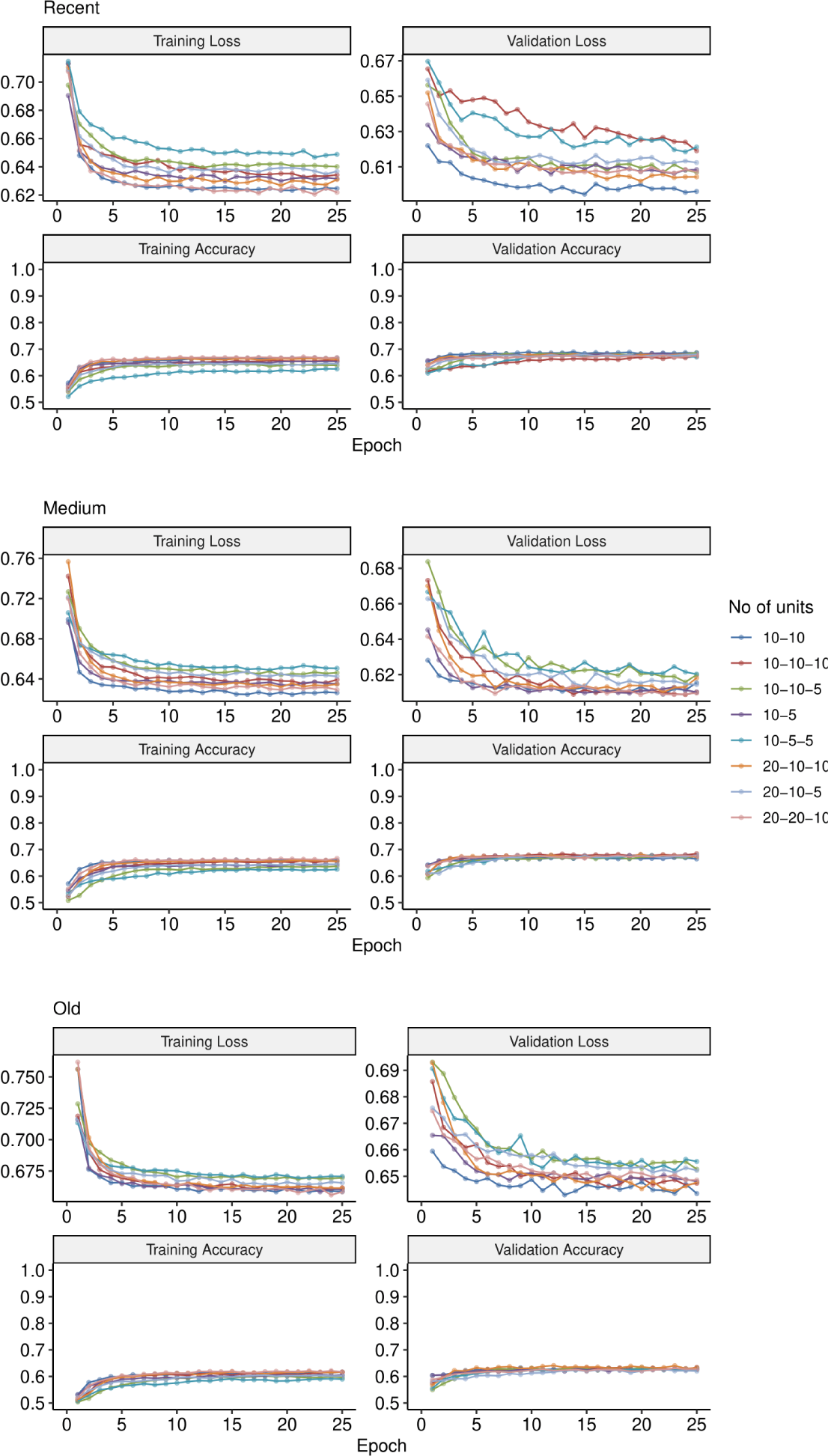
Training and validation loss and accuracy plots for hyper-parameter tuning of number of units and layers to train ANN for Test 2 (incomplete sweep *vs.* balancing selection) at different times of onset of selection (see Methods).

**Figure S15:**
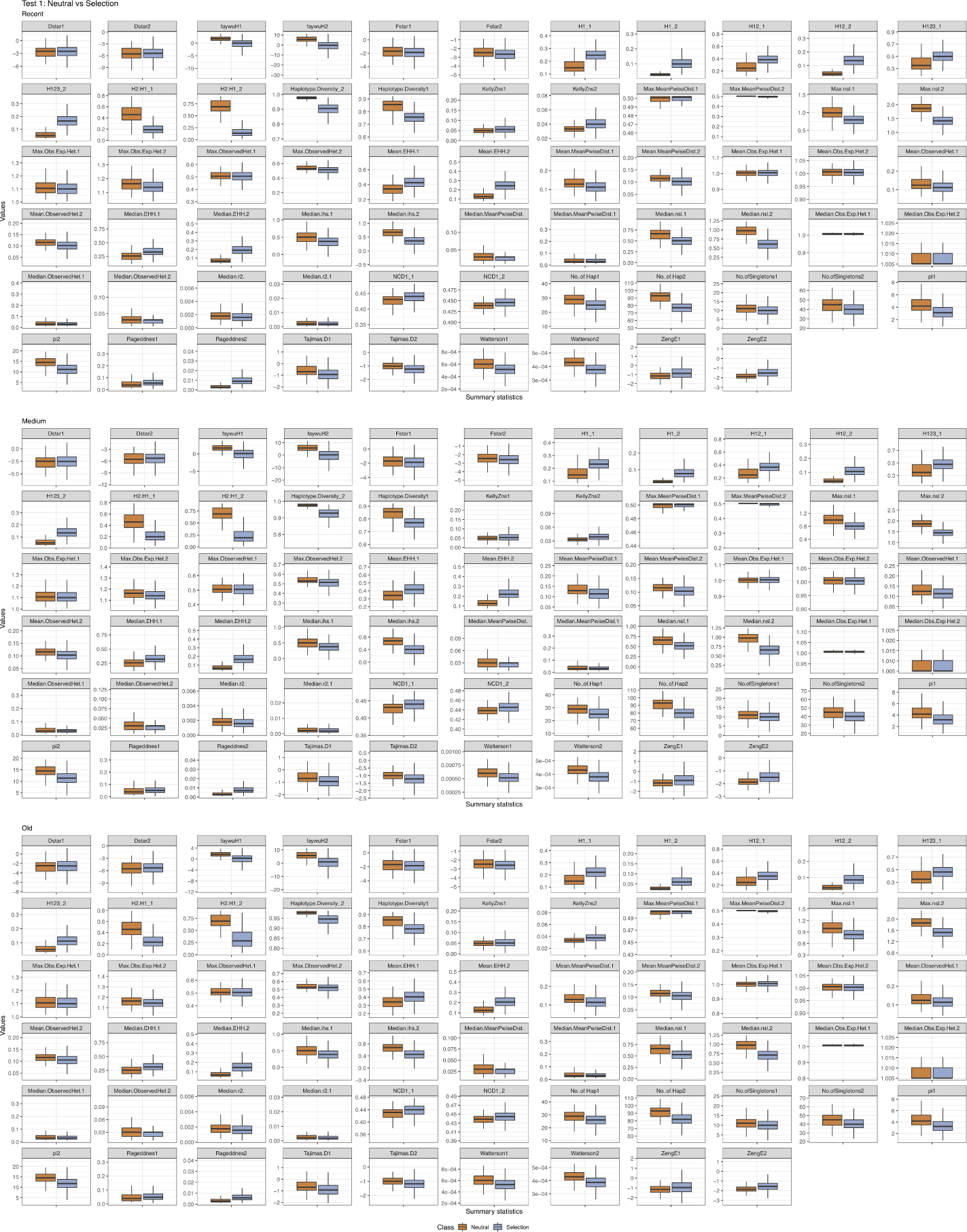
Distributions of a subset of summary statistics calculated on genes under either neutral evolution or natural selection (either ongoing positive selection or balancing selection) at different times of onset (recent, medium or old).

**Figure S16:**
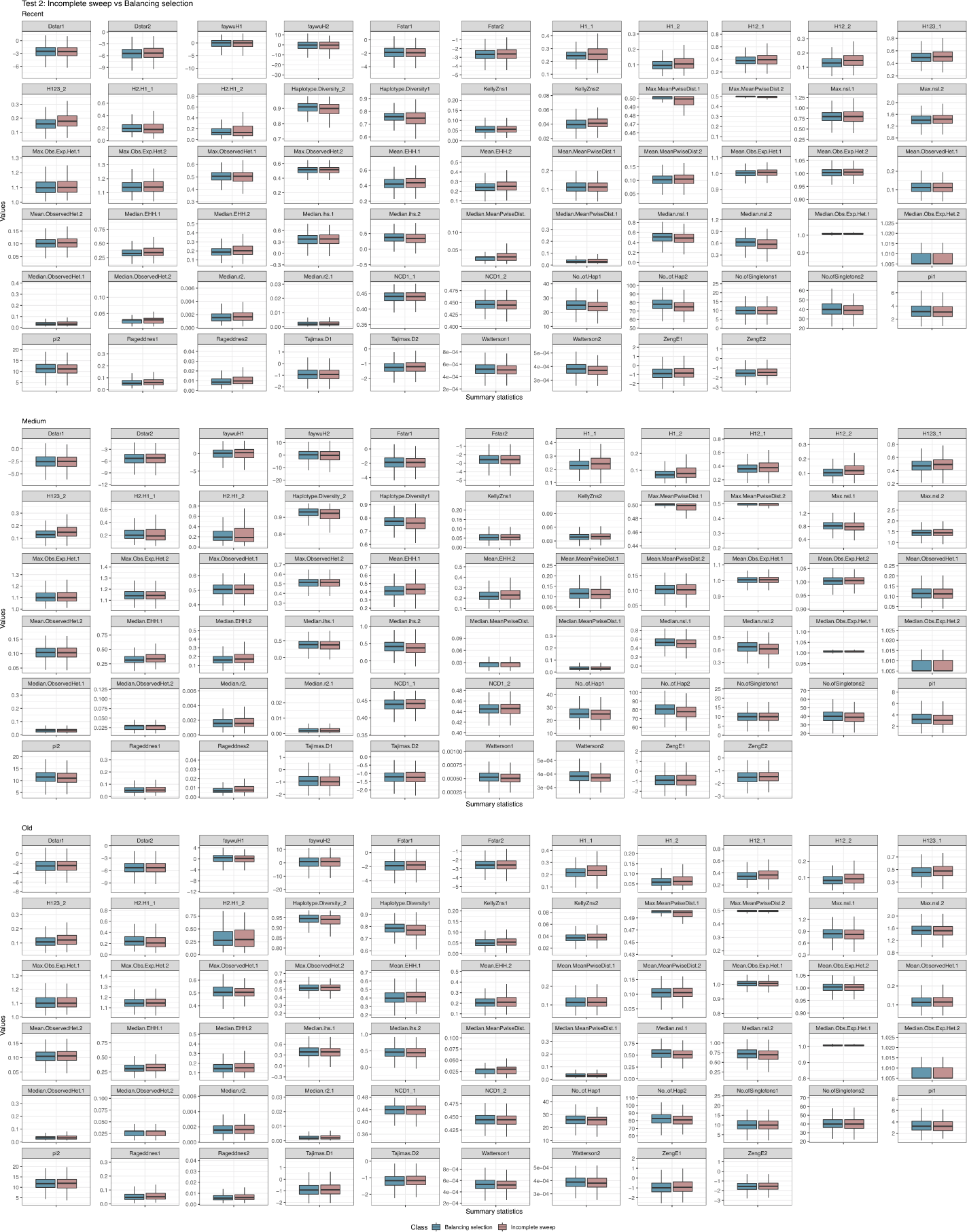
Distributions of a subset of summary statistics calculated on genes under either incomplete sweep or balancing selection at different times of onset (recent, medium or old).

**Figure S17:**
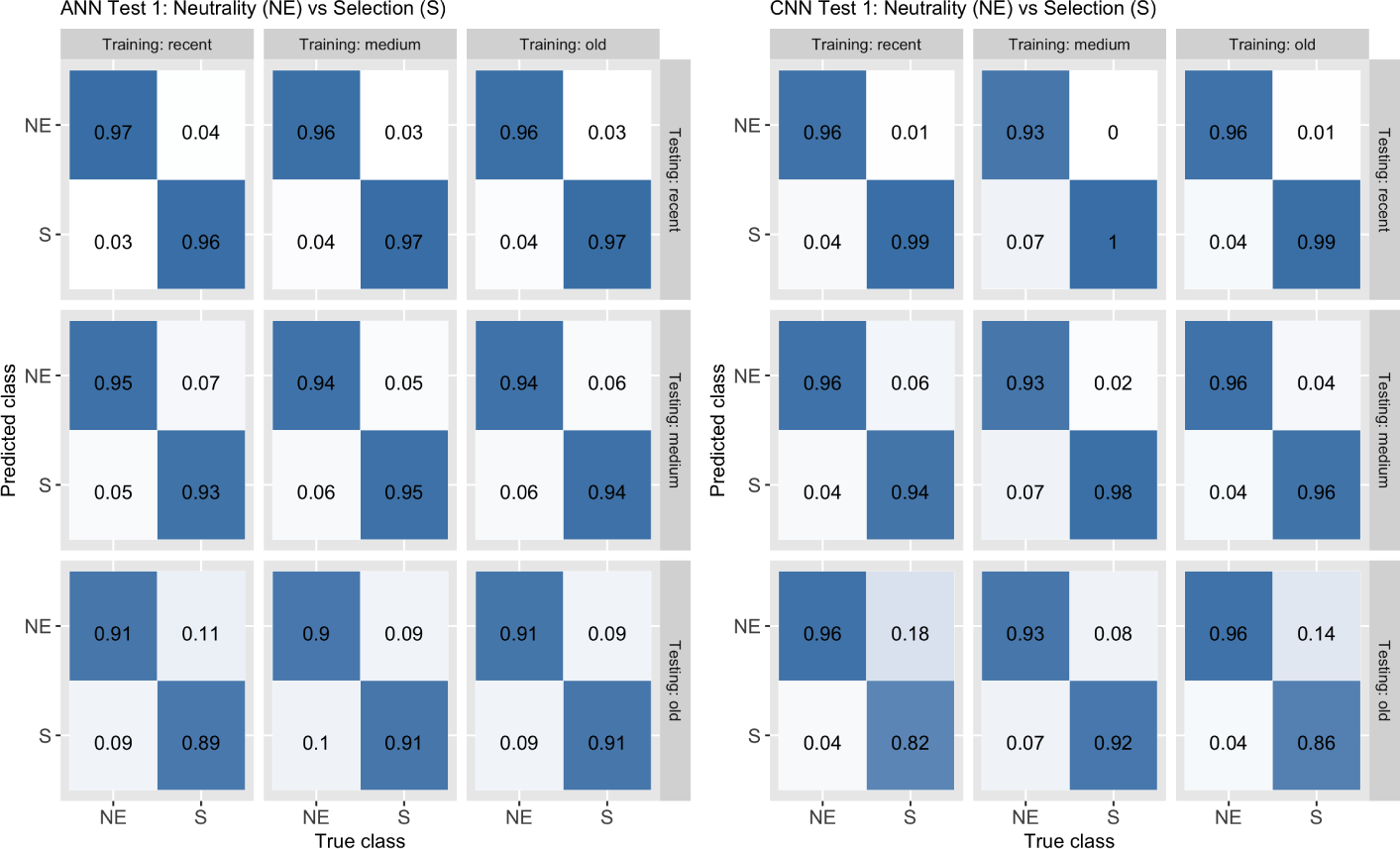
Confusion matrices accuracy for classifying loci under neutral evolution (NE) or natural selection (S) (Test 1) with both ANN and CNN for all pairs of classes for times of onset of selection (recent, medium, old) between training (y-axis) and testing data (x-axis).

**Figure S18:**
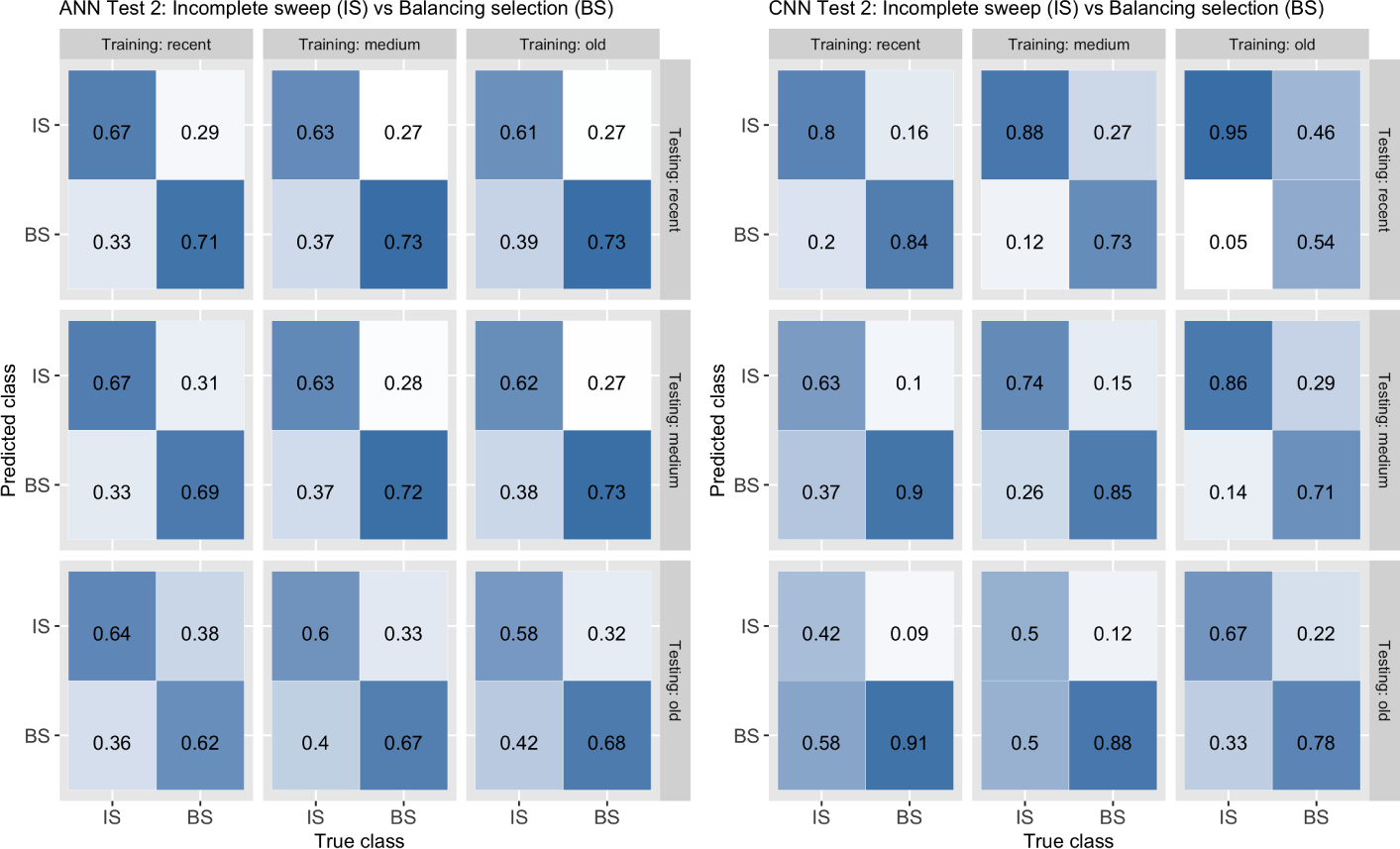
Confusion matrices accuracy for classifying loci under incomplete sweep (IS) or balancing selection (BS) (Test 2) with both ANN and CNN for all pairs of classes for times of onset of selection (recent, medium, old) between training (y-axis) and testing data (x-axis).

**Figure S19:**
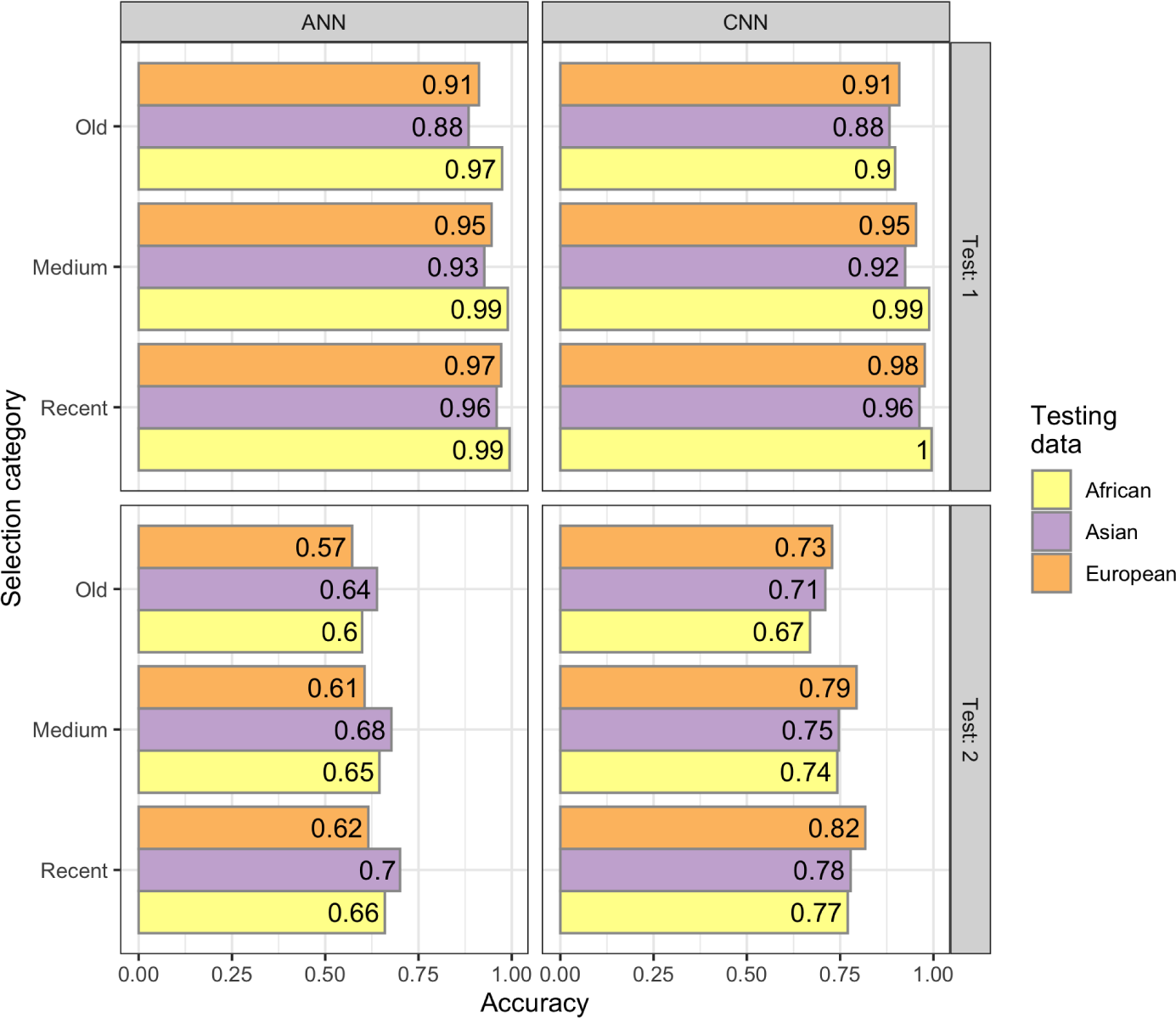
Prediction accuracy values for classifying loci under neutral evolution (NE) or natural selection (S) (Test 1), and under incomplete sweep (IS) or balancing selection (BS) (Test 2), with both ANN and CNN under a misspecified demographic model. We generated separate testing sets on different demographic models by simulating a single starting time for each selection category (23kya for recent selection, 30kya for medium selection, 37kya for old selection). Both architectures were trained using a European demographic model. As such, the label ”European” represents values where the training and testing sets where simulated from the same demographic model.

**Figure S20:**
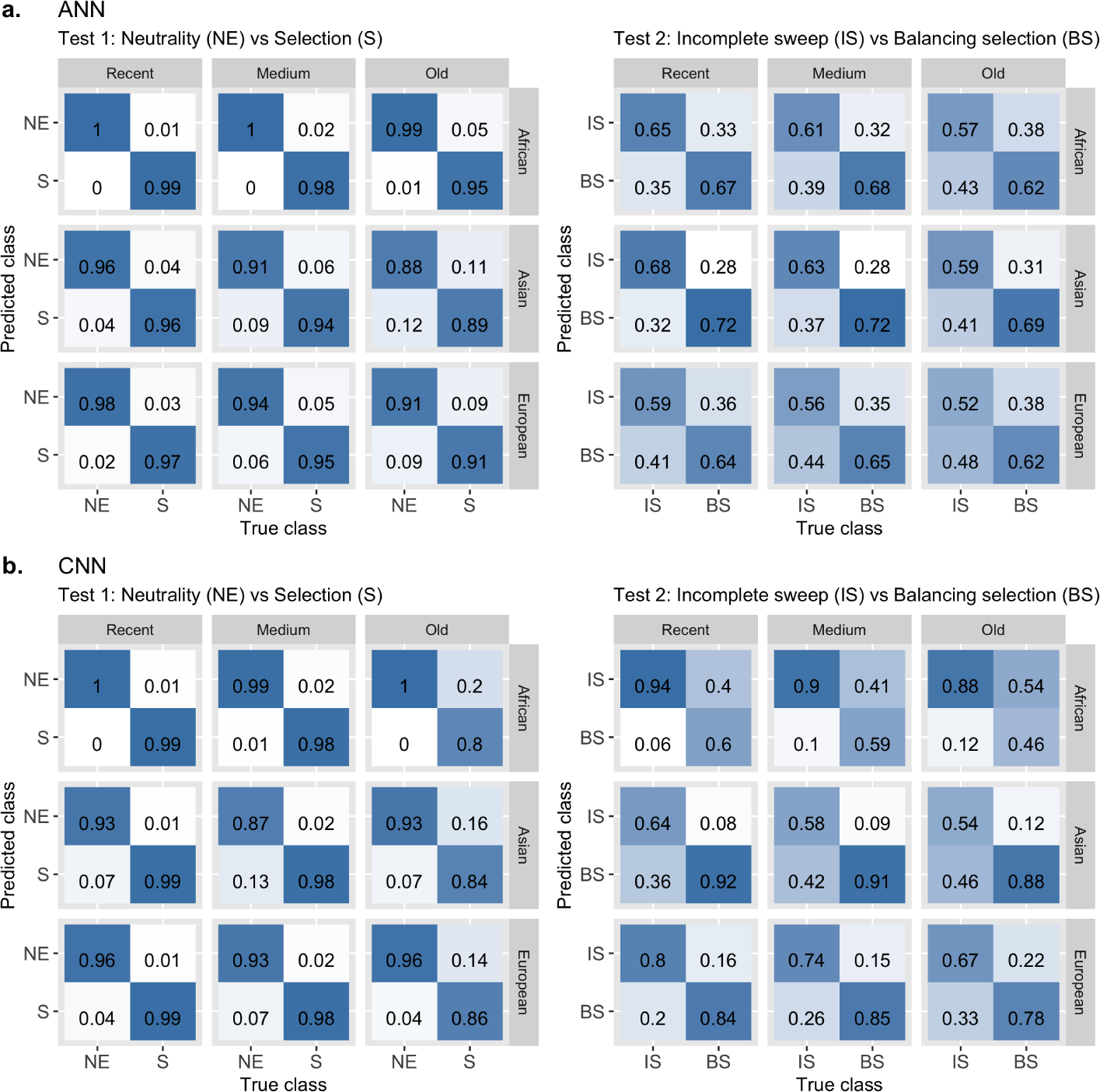
Confusion matrices for classifying loci under neutral evolution (NE) or natural selection (S) (Test 1), and under incomplete sweep (IS) or balancing selection (BS) (Test 2), with both ANN and CNN under a misspecified demographic model. We generated separate testing sets on different demographic models by simulating a single starting time for each selection category (23kya for recent selection, 30kya for medium selection, 37kya for old selection). Both architectures were trained using a European demographic model. As such, the label ”European” represents values where the training and testing sets where simulated from the same demographic model.

**Figure S21:**
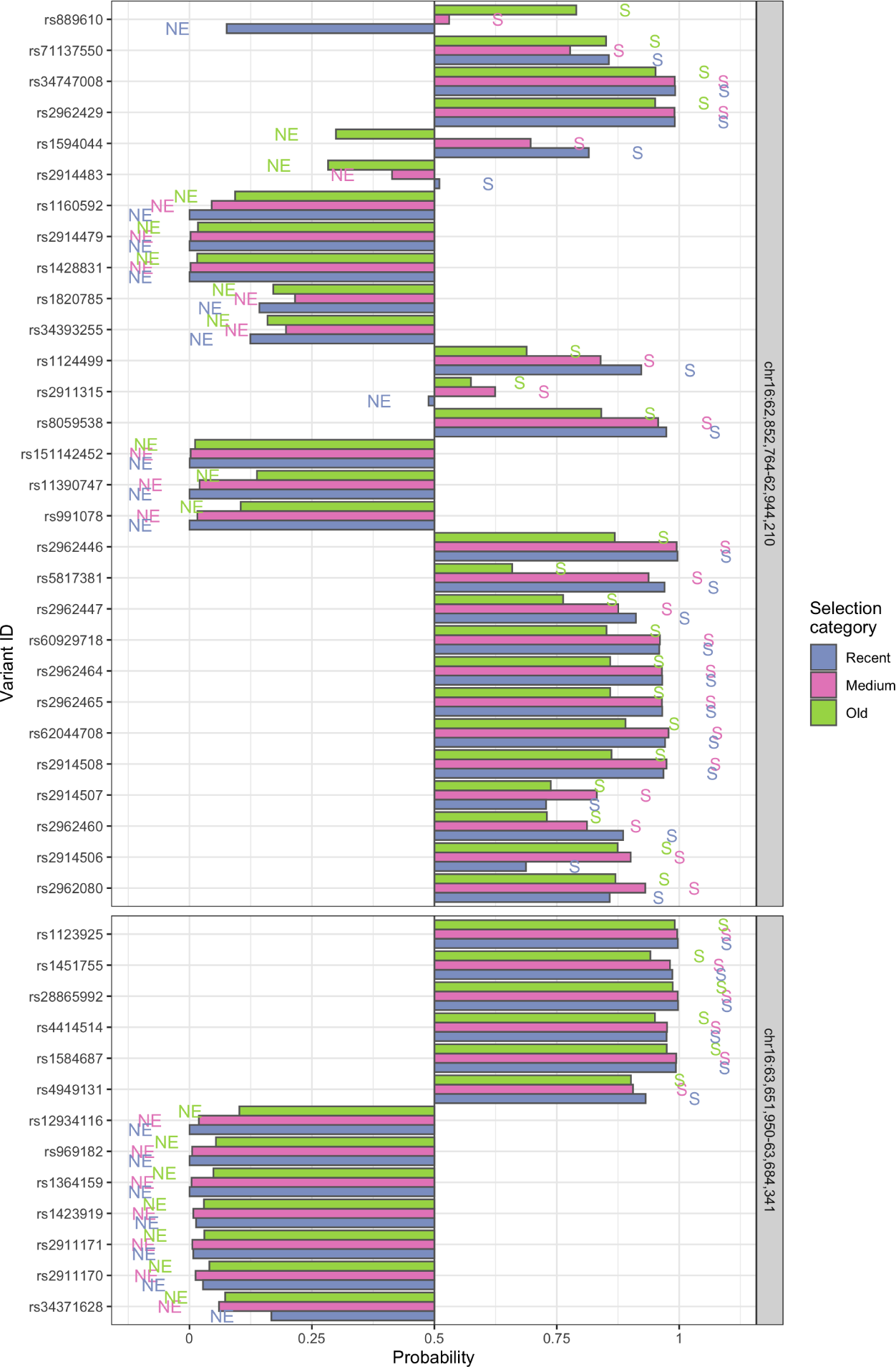
Prediction of sites under selection (S) against neutral evolution (NE) in two control neutral regions using ANN algorithm. For each site at intermediate allele frequency, the probability of being under selection (Test 1) at different times of onset (recent, medium or old) is reported.

**Figure S22:**
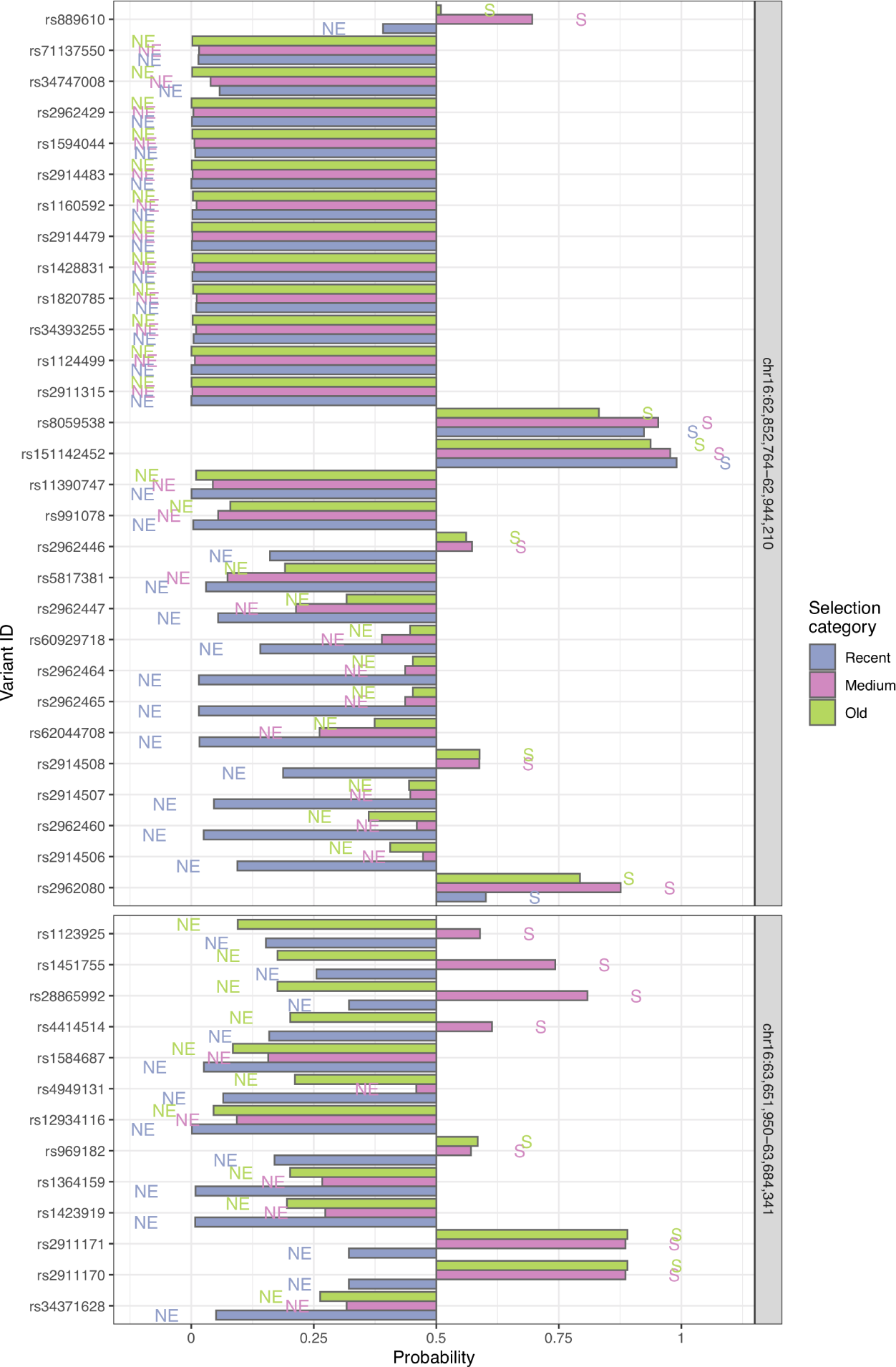
Prediction of sites under selection (S) against neutral evolution (NE) in two control neutral regions using CNN algorithm. For each site at intermediate allele frequency, the probability of being under selection (Test 1) at different times of onset (recent, medium or old) is reported.

**Figure S23:**
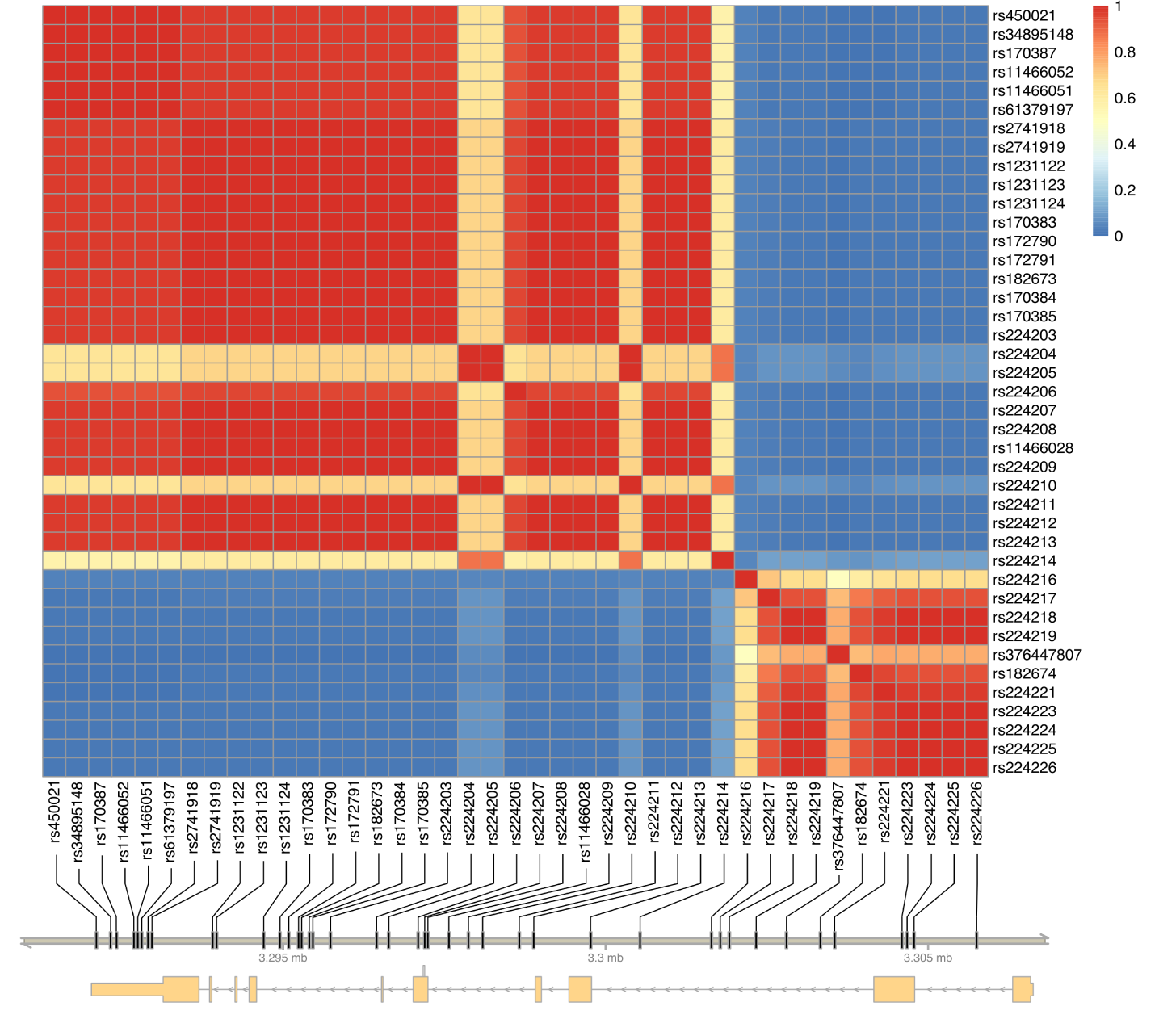
LD *r*^2^ values for all pairs of tested variants at intermediate allele frequency in the *MEFV* gene.

**Figure S24:**
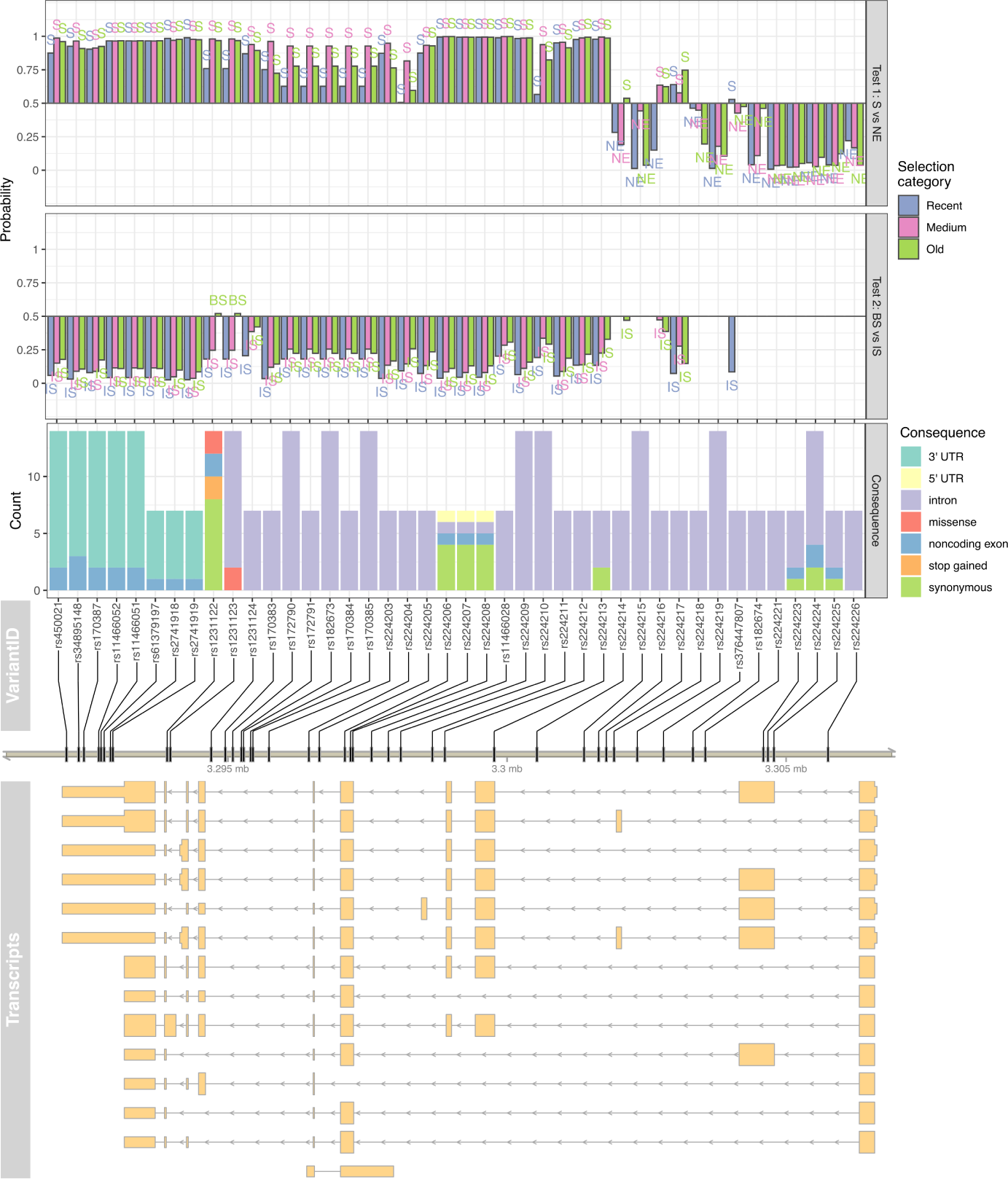
Prediction of sites under natural selection (Test 1, upper panel) or balancing selection *vs.* incomplete sweep (Test 2, second panel from top) on intermediate-frequency MEFV variants for samples from TSI population from Italy. For each tested variant, the predicted functional impact across all isoforms is reported (counts of functional consequences on third panel, genomic location on fourth panel and transcripts on fifth panel from the top).

**Figure S25:**
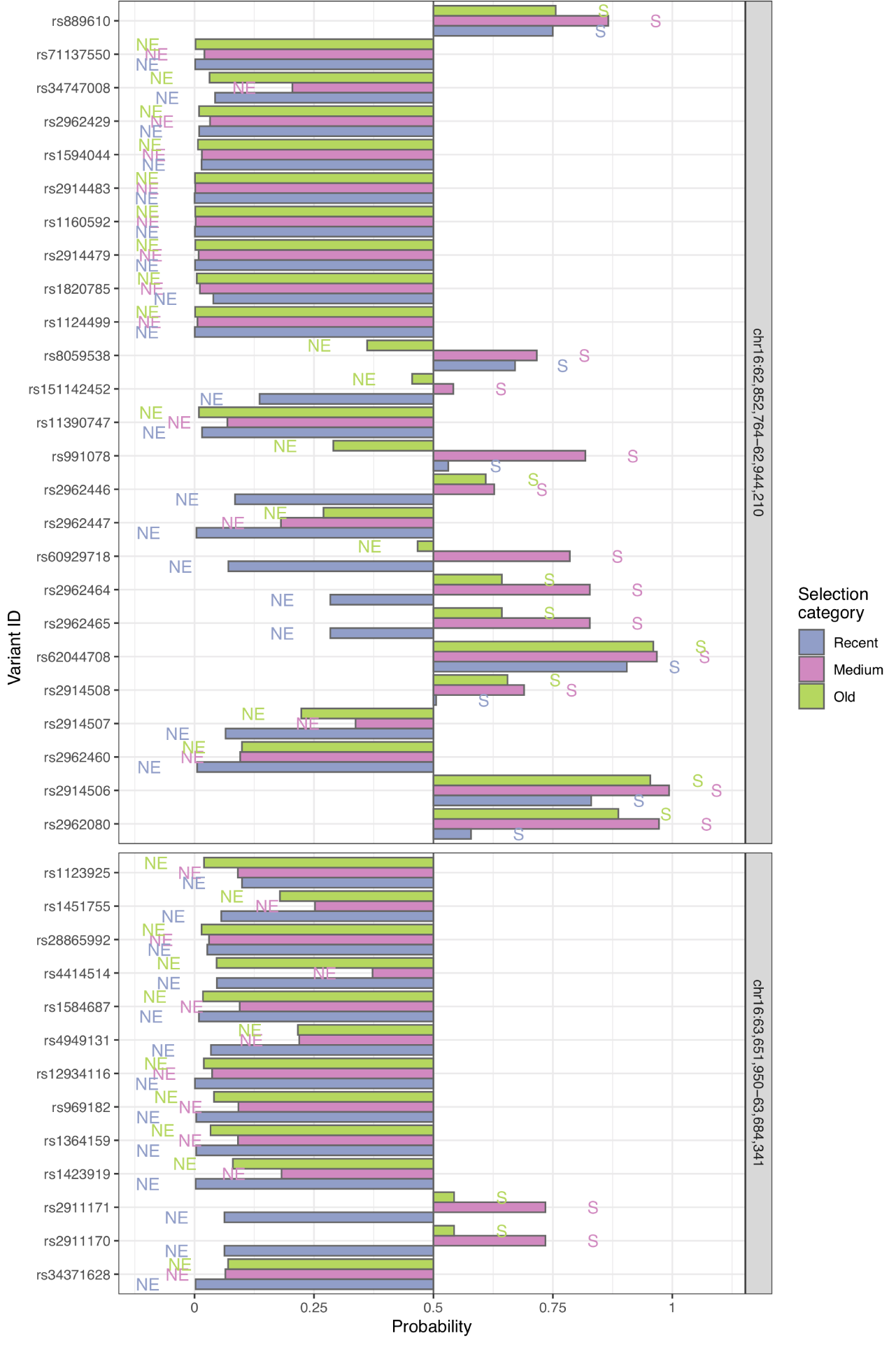
Prediction of sites under selection (S) against neutral evolution (NE) in two control neutral regions using CNN algorithm and samples from TSI population from Italy. For each site at intermediate allele frequency, the probability of being under selection (Test 1) at different time of onset (recent, medium or old) is reported.

**Figure S26:**
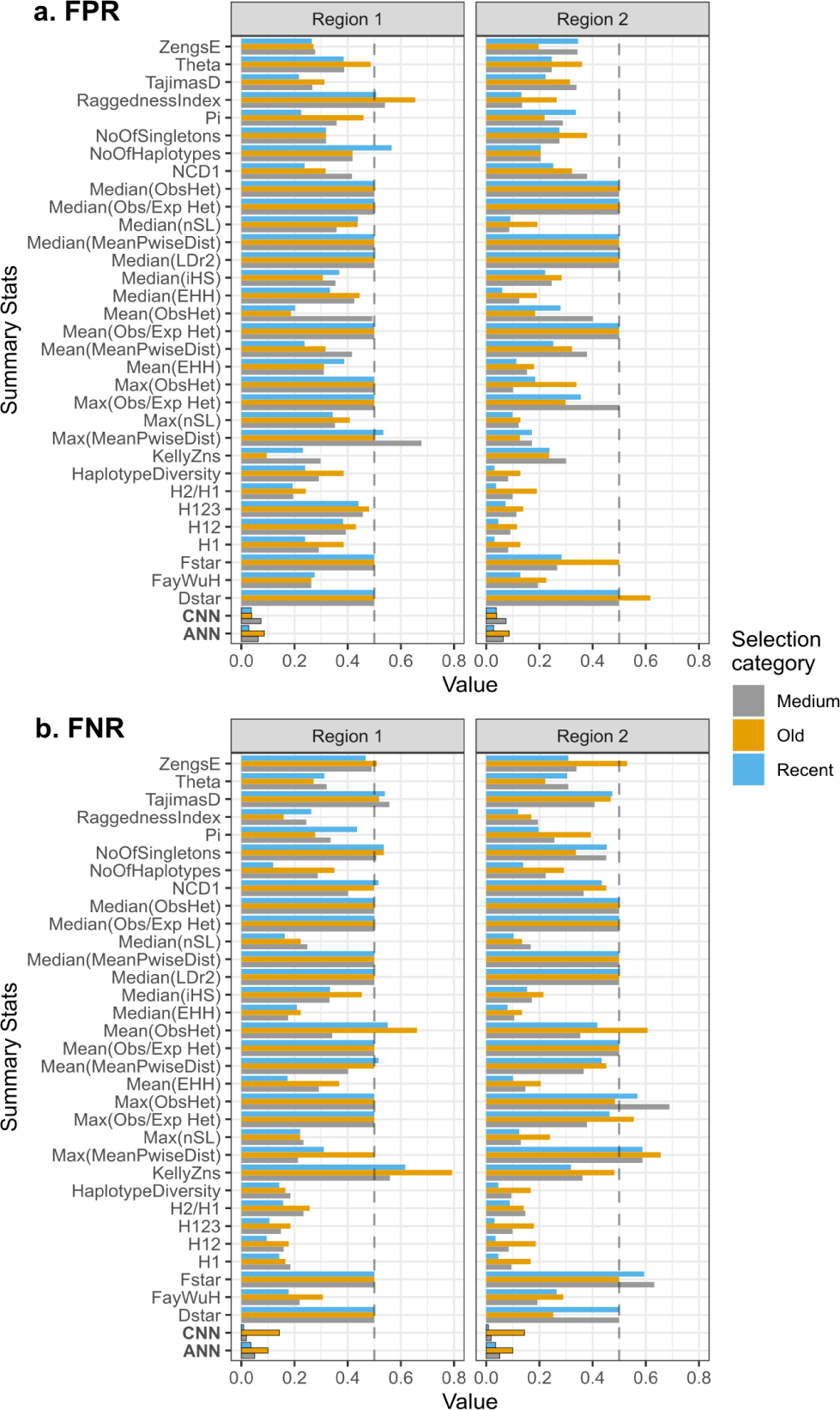
False positive (panel a) and false negative (panel b) rates for Test 1. We used cutpointr R package to find the threshold on the simulated distribution of each summary statistic that optimised the accuracy of detecting selection against neutrality. We calculated false positive and false negative rates for individual neutrality tests (where neutrality is the negative class). The y-axis shows the individual statistics, and ANN/CNN labels indicate the rates for corresponding neutral networks. Facets correspond to the two different genomic regions around the putative selected site used to calculate summary statistics. The time of selection is colour coded. The vertical dash lines sign the value of 0.5.

## References

1. Theodosius Dobzhansky. Genetics and the Origin of Species. New York: Columbia Univ. Press, 3rd editio edition, 1951.

2. Deborah Charlesworth. Balancing selection and its effects on sequences in nearby genome regions. PLoS Genetics, 2(4):379–384, 2006.

3. Saurabh Asthana, Steffen Schmidt, and Shamil Sunyaev. A limited role for balancing selection. Trends in Genetics, 21(1):30–32, 2005.

4. K L Bubb, D Bovee, D Buckley, E Haugen, M Kibukawa, M Paddock, A Palmieri, S Subramanian, Y Zhou, R Kaul, P Green, and M V Olson. Scan of human genome reveals no new loci under ancient balancing selection. Genetics, 173(4):2165–2177, aug 2006.

5. Felix M Key, Jõao C Teixeira, Cesare de Filippo, and Aida M Andŕes. Advantageous diversity maintained by balancing selection in humans. Current Opinion in Genetics and Development, 29:45–51, dec 2014.

6. Violaine Llaurens, Annabel Whibley, and Mathieu Joron. Genetic architecture and balancing selection: the life and death of differentiated variants. Molecular Ecology, 26(9):2430–2448, 2017.

7. Diogo Meyer, Richard M Single, Steven J Mack, Henry A Erlich, and Glenys Thomson. Signatures of demographic history and natural selection in the human major histocompatibility complex loci. Genetics, 173(4):2121–2142, aug 2006.

8. Anna Ferrer-Admetlla, Elena Bosch, Martin Sikora, T. Marques-Bonet, A. Ramirez-Soriano, Aura Muntasell, Arcadi Navarro, Ross Lazarus, Francesc Calafell, Jaume Bertranpetit, and Ferran Casals. Balancing Selection Is the Main Force Shaping the Evolution of Innate Immunity Genes. The Journal of Immunology, 181(2):1315–1322, jul 2008.

9. Aida M. Andŕes, Melissa J. Hubisz, Amit Indap, Dara G. Torgerson, Jeremiah D. Degenhardt, Adam R. Boyko, Ryan N. Gutenkunst, Thomas J. White, Eric D. Green, Carlos D. Bustamante, Andrew G. Clark, and Rasmus Nielsen. Targets of balancing selection in the human genome. Molecular Biology and Evolution, 26(12):2755–2764, dec 2009.

10. Michael DeGiorgio, Kirk E. Lohmueller, and Rasmus Nielsen. A Model-Based Approach for Identifying Signatures of Ancient Balancing Selection in Genetic Data. PLoS Genetics, 10(8):e1004561, aug 2014.

11. Matteo Fumagalli, Stephane M. Camus, Yoan Diekmann, Alice Burke, Marine D. Camus, Paul J. Norman, Agnel Joseph, Laurent Abi-Rached, Andrea Benazzo, Rita Rasteiro, Iain Mathieson, Maya Topf, Peter Parham, Mark G. Thomas, and Frances M. Brodsky. Genetic diversity of CHC22 clathrin impacts its function in glucose metabolism. eLife, 8, 2019.

12. Bárbara D Bitarello, Cesare de Filippo, Jõao C Teixeira, Joshua M Schmidt, Philip Kleinert, Diogo Meyer, and Aida M Andŕes. Signatures of Long-Term Balancing Selection in Human Genomes. Genome biology and evolution, 10(3):939–955, mar 2018.

13. Matteo Fumagalli, Manuela Sironi, Uberto Pozzoli, Anna Ferrer-Admettla, Linda Pattini, and Rasmus Nielsen. Signatures of environmental genetic adaptation pinpoint pathogens as the main selective pressure through human evolution. PLoS Genetics, 7(11):e1002355, nov 2011.

14. Rachele Cagliani, Matteo Fumagalli, Stefania Riva, Uberto Pozzoli, Giacomo P. Comi, Nereo Bresolin, and Manuela Sironi. Genetic variability in the ACE gene region surrounding the Alu I/D polymorphism is maintained by balancing selection in human populations. Pharmacogenetics and Genomics, 20(2):131–134, 2010.

15. Joris R. Delanghe, Marijn M. Speeckaert, and Marc L. De Buyzere. COVID-19 infections are also affected by human ACE1 D/I polymorphism. Clinical chemistry and laboratory medicine, pages 1–2, 2020.

16. Anna Fijarczyk and Wies-law Babik. Detecting balancing selection in genomes: limits and prospects. Molecular Ecology, 24(14):3529–3545, jul 2015.

17. M. Fumagalli, R. Cagliani, U. Pozzoli, S. Riva, G. P. Comi, G. Menozzi, N. Bresolin, and M. Sironi. A population genetics study of the familial mediterranean fever gene: Evidence of balancing selection under an over-dominance regime. Genes and Immunity, 10(8):678–686, 2009.

18. Rachele Cagliani, Matteo Fumagalli, Stefania Riva, Uberto Pozzoli, Marco Fracassetti, Nereo Bresolin, Giacomo P. Comi, and Manuela Sironi. Polymorphisms in the CPB2 gene are maintained by balancing selection and result in haplotype-preferential splicing of exon 7. Molecular Biology and Evolution, 27(8):1945–1954, 2010.

19. Vivak Soni, Michiel Vos, and Adam Eyre-Walker. A new test suggests that balancing selection maintains hundreds of non-synonymous polymorphisms in the human genome. bioRxiv, 2021.

20. Katherine M. Siewert and Benjamin F. Voight. Detecting long-term balancing selection using allele frequency correlation. Molecular Biology and Evolution, 34(11):2996–3005, nov 2017.

21. Rachele Cagliani, Matteo Fumagalli, Stefania Riva, Uberto Pozzoli, Giacomo P. Comi, Giorgia Menozzi, Nereo Bresolin, and Manuela Sironi. The signature of long-standing balancing selection at the human defensin *β*-1 promoter. Genome Biology, 9(9), 2008.

22. Ellen M Leffler, Ziyue Gao, Susanne Pfeifer, Laure Śegurel, Adam Auton, Oliver Venn, Rory Bowden, Ronald Bontrop, Jeffrey D Wall, Guy Sella, Peter Donnelly, Gilean McVean, and Molly Przeworski. Multiple instances of ancient balancing selection shared between humans and chimpanzees. Science, 340(6127):1578–1582, mar 2013.

23. Jõao C. Teixeira, Cesare De Filippo, Antje Weihmann, Juan R. Meneu, Fernando Racimo, Michael Dannemann, Birgit Nickel, Anne Fischer, Michel Halbwax, Claudine Andre, Rebeca Atencia, Matthias Meyer, Geńıs Parra, Svante Pääbo, and Aida M. Andŕes. Long-term balancing selection in LAD1 maintains a missense trans-species polymorphism in humans, chimpanzees, and bonobos. Molecular Biology and Evolution, 32(5):1186–1196, 2015.

24. Xiaoheng Cheng and Michael DeGiorgio. Flexible mixture model approaches that accommodate footprint size variability for robust detection of balancing selection. Molecular Biology and Evolution, pages 1–40, 2020.

25. Matteo Fumagalli, Rachele Cagliani, Uberto Pozzoli, Stefania Riva, Giacomo P G.P. Comi, Giorgia Menozzi, Nereo Bresolin, and Manuela Sironi. Widespread balancing selection and pathogen-driven selection at blood group antigen genes. Genome research, 19(2):199–212, feb 2009.

26. Diamantis Sellis, Benjamin J. Callahan, Dmitri A. Petrov, and Philipp W. Messer. Heterozygote advantage as a natural consequence of adaptation in diploids. Proceedings of the National Academy of Sciences of the United States of America, 108(51):20666–20671, 2011.

27. Daniel R. Schrider and Andrew D. Kern. Supervised Machine Learning for Population Genetics: A New Paradigm. Trends in Genetics, 34(4):301–312, apr 2018.

28. Mehreen R. Mughal and Michael DeGiorgio. Localizing and classifying adaptive targets with trend filtered regression. Molecular Biology and Evolution, 36(2):252–270, 2019.

29. Daniel R. Schrider and Andrew D. Kern. S/HIC: Robust Identification of Soft and Hard Sweeps Using Machine Learning. PLoS Genetics, 12(3):1–31, 2016.

30. Andrew D. Kern and Daniel R. Schrider. DiploS/HIC: An updated approach to classifying selective sweeps. G3: Genes, Genomes, Genetics, 8(6):1959–1970, 2018.

31. Lauren Alpert Sugden, Elizabeth G. Atkinson, Annie P. Fischer, Stephen Rong, Brenna M. Henn, and Sohini Ramachandran. Localization of adaptive variants in human genomes using averaged one-dependence estimation. Nature Communications, 9(1), 2018.

32. Pavlos Pavlidis, Jeffrey D. Jensen, and Wolfgang Stephan. Searching for footprints of positive selection in whole-genome SNP data from nonequilibrium populations. Genetics, 185(3):907–922, 2010.

33. Roy Ronen, Nitin Udpa, Eran Halperin, and Vineet Bafna. Learning natural selection from the site frequency spectrum. Genetics, 195(1):181–193, 2013.

34. Kao Lin, Haipeng Li, Christian Schlötterer, and Andreas Futschik. Distinguishing positive selection from neutral evolution: Boosting the performance of summary statistics. Genetics, 187(1):229–244, 2011.

35. Yann Lecun, Yoshua Bengio, and Geoffrey Hinton. Deep learning. Nature, 521(7553):436–444, 2015.

36. Sara Sheehan and Yun S. Song. Deep Learning for Population Genetic Inference. PLoS Computational Biology, 12(3):e1004845, mar 2016.

37. Alex Krizhevsky, Ilya SutskeverI, and Geoffrey Hinton. ImageNet Classification with Deep ConvolutionalNeural Networks. Advances in neural information processing systems, pages 1097–1105, 2012.

38. Lex Flagel, Yaniv Brandvain, and Daniel R Schrider. The Unreasonable Effectiveness of Convolutional Neural Networks in Population Genetic Inference. Molecular biology and evolution, 36(2):220–238, dec 2019.

39. Jeffrey Chan, Jeffrey P. Spence, Sara Mathieson, Valerio Perrone, Paul A. Jenkins, and Yun S. Song. A likelihood-free inference framework for population genetic data using exchangeable neural networks. Advances in Neural Information Processing Systems, 2018-December(NeurIPS 2018):8594–8605, 2018.

40. Luis Torada, Lucrezia Lorenzon, Alice Beddis, Ulas Isildak, Linda Pattini, Sara Mathieson, and Matteo Fumagalli. ImaGene: a convolutional neural network to quantify natural selection from genomic data. BMC Bioinformatics, 20(S9):337, nov 2019.

41. Théophile Sanchez, Jean Cury, Guillaume Charpiat, and Flora Jay. Deep learning for population size history inference: Design, comparison and combination with approximate Bayesian computation. Molecular Ecology Resources, 00(July):1–16, 2020.

42. Alexander T Xue, Daniel R Schrider, Andrew D Kern, and Ag1000g Consortium. Discovery of Ongoing Selective Sweeps within Anopheles Mosquito Populations Using Deep Learning. Molecular Biology and Evolution, 10 2020. msaa259.

43. Y. Lecun, L. Bottou, Y. Bengio, and P. Haffner. Gradient-based learning applied to document recognition. Proceedings of the IEEE, 86(11):2278–2324, 1998.

44. Jiuxiang Gu, Zhenhua Wang, Jason Kuen, Lianyang Ma, Amir Shahroudy, Bing Shuai, Ting Liu, Xingxing Wang, Gang Wang, Jianfei Cai, and Tsuhan Chen. Recent advances in convolutional neural networks. Pattern Recognition, 77:354–377, may 2018.

45. I. Touitou. The spectrum of Familial Mediterranean Fever (FMF) mutations. European Journal of Human Genetics, 9(7):473–483, 2001.

46. Yong Hwan Park, Elaine F. Remmers, Wonyong Lee, Amanda K. Ombrello, Lawton K. Chung, Zhao Shilei, Deborah L. Stone, Maya I. Ivanov, Nicole A. Loeven, Karyl S. Barron, Patrycja Hoffmann, Michele Nehrebecky, Yeliz Z. Akkaya-Ulum, Erdal Sag, Banu Balci-Peynircioglu, Ivona Aksentijevich, Ahmet Gül, Charles N. Rotimi, Hua Chen, James B. Bliska, Seza Ozen, Daniel L. Kastner, Daniel Shriner, and Jae Jin Chae. Ancient familial Mediterranean fever mutations in human pyrin and resistance to Yersinia pestis. Nature Immunology, 2020.

47. Benjamin C Haller and Philipp W Messer. SLiM 3: Forward Genetic Simulations Beyond the Wright-Fisher Model. Molecular Biology and Evolution, 36(3):632–637, mar 2019.

48. Julien Jouganous, Will Long, Aaron P. Ragsdale, and Simon Gravel. Inferring the joint demographic history of multiple populations: Beyond the diffusion approximation. Genetics, 206(3):1549–1567, jul 2017.

49. 1000 Genomes Project Consortium. A global reference for human genetic variation. Nature, 526(7571):68–74, oct 2015.

50. Benjamin M. Peter, Emilia Huerta-Sanchez, and Rasmus Nielsen. Distinguishing between Selective Sweeps from Standing Variation and from a De Novo Mutation. PLoS Genetics, 8(10), 2012.

51. M. Nei and W. H. Li. Mathematical model for studying genetic variation in terms of restriction endonucleases. Proceedings of the National Academy of Sciences of the United States of America, 76(10):5269–5273, 1979.

52. G. A. Watterson. On the number of segregating sites in genetical models without recombination. Theoretical Population Biology, 7(2):256–276, apr 1975.

53. Fumio Tajima. Statistical analysis of DNA polymorphism. Japanese Journal of Genetics, 68(6):567–595, dec 1993.

54. W. G. Hill and Alan Robertson. Linkage disequilibrium in finite populations. Theoretical and Applied Genetics, 38(6):226–231, jun 1968.

55. John K. Kelly. A test of neutrality based on interlocus associations. Genetics, 146(3):1197–1206, 1997.

56. Y X Fu and W H Li. Statistical tests of neutrality of mutations. Genetics, 133(3):693–709, 1993.

57. Nandita R. Garud, Philipp W. Messer, Erkan O. Buzbas, and Dmitri A. Petrov. Recent Selective Sweeps in North American Drosophila melanogaster Show Signatures of Soft Sweeps. PLoS Genetics, 11(2):1–32, feb 2015.

58. Benjamin F. Voight, Sridhar Kudaravalli, Xiaoquan Wen, and Jonathan K. Pritchard. A map of recent positive selection in the human genome. PLoS Biology, 4(3):0446–0458, 2006.

59. Pardis C. Sabeti, David E. Reich, John M. Higgins, Haninah Z.P. Levine, Daniel J. Richter, Stephen F. Schaffner, Stacey B. Gabriel, Jill V. Platko, Nick J. Patterson, Gavin J. McDonald, Hans C. Ackerman, Sarah J. Camp-bell, David Altshuler, Richard Cooper, Dominic Kwiatkowski, Ryk Ward, and Eric S. Lander. Detecting recent positive selection in the human genome from haplotype structure. Nature, 419(6909):832–837, oct 2002.

60. Kai Zeng, Yun Xin Fu, Suhua Shi, and Chung I. Wu. Statistical tests for detecting positive selection by utilizing high-frequency variants. Genetics, 174(3):1431–1439, nov 2006.

61. Justin C Fay and Chung I. Wu. Hitchhiking under positive Darwinian selection. Genetics, 155(3):1405–1413, 2000.

62. Anna Ferrer-Admetlla, Mason Liang, Thorfinn Korneliussen, and Rasmus Nielsen. On detecting incomplete soft or hard selective sweeps using haplotype structure. Molecular Biology and Evolution, 31(5):1275–1291, 2014.

63. H.C. Harpending. Signature of Ancient Population Growth in a Low-Resolution Mitochondrial DNA Mismatch Distribution. Human Biology, 66(4):591–600, 1994.

64. F. Pedregosa, G. Varoquaux, A. Gramfort, V. Michel, B. Thirion, O. Grisel, M. Blondel, P. Prettenhofer, R. Weiss, V. Dubourg, J. Vanderplas, A. Passos, D. Cournapeau, M. Brucher, M. Perrot, and E. Duchesnay. Scikit-learn: Machine learning in Python. Journal of Machine Learning Research, 12:2825–2830, 2011.

65. Francois Chollet et al. Keras. https://keras.io, 2015.

66. Diederik P. Kingma and Jimmy BaAdam: A method for stochastic optimization, 2014.

67. Ruder S. An overview of gradient descent optimization algorithms, 2017.

68. Johannes Rainer. EnsDb.Hsapiens.v75: Ensembl based annotation package, 2017. R package version 2.99.0.

69. Florian Hahne and Robert Ivanek. *Statistical Genomics: Methods and Protocols*, chapter Visualizing Genomic Data Using Gviz and Bioconductor, pages 335–351. Springer New York, New York, NY, 2016.

70. Leonardo Arbiza, Elaine Zhong, and Alon Keinan. NRE: A tool for exploring neutral loci in the human genome. BMC Bioinformatics, 13(1):1, 2012.

71. Hadley Wickham. ggplot2: Elegant Graphics for Data Analysis. Springer-Verlag New York, 2016.

72. Alboukadel Kassambara. ggpubr: ’ggplot2’ Based Publication Ready Plots, 2020. R package version 0.3.0.

73. Raivo Kolde. pheatmap: Pretty Heatmaps, 2018. R package version 1.0.12.

74. Linlin Shen and Li Bai. A review on Gabor wavelets for face recognition. Pattern Analysis and Applications, 9(2-3):273–292, 2006.

75. David G. Lowe. Object recognition from local scale-invariant features. Proceedings of the IEEE International Conference on Computer Vision, 2:1150–1157, 1999.

76. John M. Gauch. Image segmentation and analysis via multiscale gradient watershed hierarchies. IEEE Transactions on Image Processing, 8(1):69–79, 1999.

77. Dzmitry Bahdanau, Kyung Hyun Cho, and Yoshua Bengio. Neural machine translation by jointly learning to align and translate. 3rd International Conference on Learning Representations, ICLR 2015 - Conference Track Proceedings, pages 1–15, 2015.

78. Martin Wistuba, Ambrish Rawat, and Tejaswini Pedapati. A Survey on Neural Architecture Search. 2019.

79. Thorfinn Sand Korneliussen, Ida Moltke, Anders Albrechtsen, and Rasmus Nielsen. Calculation of Tajima’s D and other neutrality test statistics from low depth next-generation sequencing data. BMC Bioinformatics, 14(1), 2013.

80. Philip Schaner, Neil Richards, Anish Wadhwa, Ivona Aksentijevich, Daniel Kastner, Priscilla Tucker, and Deborah Gumucio. Episodic evolution of pyrin in primates: Human mutations recapitulate ancestral amino acid states. Nature Genetics, 27(3):318–321, 2001.

81. Oskar Schnappauf, Jae Jin Chae, Daniel L. Kastner, and Ivona Aksentijevich. The Pyrin Inflammasome in Health and Disease. Frontiers in immunology, 10(August):1745, 2019.

82. Je Wook Yu, Teresa Fernandes-Alnemri, Pinaki Datta, Jianghong Wu, Christine Juliana, Leobaldo Solorzano, Margaret McCormick, Zhi Jia Zhang, and Emad S. Alnemri. Pyrin Activates the ASC Pyroptosome in Response to Engagement by Autoinflammatory PSTPIP1 Mutants. Molecular Cell, 28(2):214–227, 2007.

83. Alessandro Stella, Fabiana Cortellessa, Giuseppe Scaccianoce, Barbara Pivetta, Enrica Settimo, and Piero Portincasa. Familial Mediterranean fever: Breaking all the (genetic) rules. Rheumatology (United Kingdom), 58(3):463–467, 2019.

84. Matteo Accetturo, Angela Maria D’Uggento, Piero Portincasa, and Alessandro Stella. Improvement of MEFV gene variants classification to aid treatment decision making in familial Mediterranean fever. Rheumatology (United Kingdom), 59(4):754–761, 2020.

85. Marianne Dehasque, María C. Ávila-Arcos, David Díez-del-Molino, Matteo Fumagalli, Katerina Guschanski, Eline D. Lorenzen, Anna-Sapfo Malaspinas, Tomas Marques-Bonet, Michael D. Martin, Gemma G. R. Murray, Alexander S. T. Papadopulos, Nina Overgaard Therkildsen, Daniel Weg-mann, Love Daĺen, and Andrew D. Foote. Inference of natural selection from ancient DNA. Evolution Letters, 4(2):94–108, 2020.

86. Etienne Patin. Plague as a cause for familial Mediterranean fever. Nature Immunology, pages 4–5, 2020.

87. Simon Gravel, Brenna M. Henn, Ryan N. Gutenkunst, Amit R. Indap, Gabor T. Marth, Andrew G. Clark, Fuli Yu, Richard A. Gibbs, and Carlos D. Bustamante. Demographic history and rare allele sharing among human populations. Proceedings of the National Academy of Sciences, 108(29):11983–11988, 2011.

